# Immunopeptidomics-guided identification of functional neoantigens in non-small cell lung cancer

**DOI:** 10.1101/2024.05.30.596609

**Authors:** Ben Nicholas, Alistair Bailey, Katy J McCann, Oliver Wood, Eve Currall, Peter Johnson, Tim Elliott, Christian Ottensmeier, Paul Skipp

## Abstract

Non-small cell lung cancer (NSCLC) has poor survival even with modern checkpoint inhibitor therapies. Personalised vaccines based on short peptide neoantigens containing tumour mutations are an attractive precision medicine strategy, but identifying therapeutically relevant neoantigens remains challenging, with existing methods yielding positive responses in only 6% of candidates tested.

We developed an immunopeptidomics approach to improve neoantigen identification in 24 NSCLC patients (15 adenocarcinoma, 9 squamous cell carcinoma). We directly identified one neoantigen and using whole exome sequencing, transcriptomics and mass spectrometry-based immunopeptidomics, we filtered predicted neoantigens based on observed cohort HLA peptide presentation. This approach achieved positive functional responses in 5 of 6 patients tested (83% success rate) with 13% of putative neoantigens (9 out of 70) eliciting strong responses. Bayesian modelling of our initial rules-based neoantigen selection further revealed patient specific peptide presentation patterns and propensities.

Our findings demonstrate that incorporating donor specific HLA peptide presentation data substantially improves neoantigen identification success rates and immune response specificity, advancing personalised cancer vaccine development.

## Introduction

Lung cancer is the second most common cancer in the UK and despite recent advances in checkpoint inhibitor therapies fewer than 20% of patients survive 5 years, with most surviving less than one year post-diagnosis^1,2^.The majority are diagnosed at advanced stages (III-IV) with non-small cell lung cancer (NSCLC) accounting for 85-90% of cases. NSCLC comprises three histological subtypes: adenocarcinoma (LUAD, most common, peripheral), squamous cell carcinoma (LUSC, central), and large cell undifferentiated carcinoma (least common, can form throughout the lung).

Four immunotherapies targeting PD-1/PD-L1 are licensed for NSCLC but show reduced efficacy in patients with EGFR or ALK mutations^3,4^. Consequently, immunotherapy typically follows chemotherapy and/or targeted therapy. While LUAD generally shows more favourable survival than LUSC, treatment resistance remains a major challenge.

The TRACERx study revealed evolutionary processes underlying this resistance. Whole genome doubling, driven by tobacco smoke and cytidine deaminase activity, protects tumours against high mutation burdens and chromosomal instability^5^. Smoking-induced truncal mutations and deaminase-driven branch mutations create extensive intratumour heterogeneity that predicts recurrence and death, yet confounds biomarker utility for immunotherapy response prediction^6^.

An attractive strategy for NSCLC treatment is vaccination targeting on HLA presented neoantigens. This approach assumes neoantigens can be identified that expand tumour killing T-cell populations and/or modulate the tumour microenvironment to make T-cell infiltration or checkpoint inhibitors more effective. The personalised nature of neoantigens minimise the risk of off-target effects and autoimmunity. However, direct identification of neoantigens is rare^7,8^ and most approaches to neoantigen discovery rely on predicting that a given mutation leads to protein synthesis, antigen processing and HLA presentation. Direct observation is rare in part due to limits in the sensitivity of the mass spectrometry proteomic detection of HLA ligands, known as immunopeptidomics. Moreover, it is estimated that only a small fraction of mutations are actually presented, possibly as low as 0.5% of non-silent mutations^9^. For example, a NSCLC tumour with 600 missense variants might yield only 3 presented neoantigens amongst a lung tissue immunopeptidome of around 60,000 unique class I and II HLA peptides^10^. A typical experiment may identify 3,000 of these peptides. Assuming a hypergeometric distribution, the probability of observing one class I or II HLA neoantigen is about 14%. Or to put it another way, there is around an 86% chance of not seeing any neoantigens in any single mass spectrometry proteomics experiment.

Given these odds, much effort has been put into the in silico prediction of mutations that will give rise to neoantigens that would make effective vaccines. There are well established algorithms that can predict the likelihood of a peptide of given amino acid sequence binding to an HLA molecule, and immunopeptidomic evidence of the peptide length preference of peptide for different HLA allotypes^11^, and preferential regions of proteins favourable for presentation^12–14^. However, even with this knowledge prediction is stymied by the number of potential neoantigen candidates each mutation might yield, creating large lists of candidate peptides. Moreover, the key biochemical and structural parameters of immunogenic neoantigens remain unknown. The best neoantigen prediction models have a success rate such that around 6% of their putative neoantigens are T-cell reactive^15,16^, although recent machine learning models report increased predictive power^17–20^.

Here we adopted a patient-specific approach where, rather than relying solely on various general HLA peptide characteristics, we leveraged each patient’s HLA-I and -II immunopeptidomes alongside the cohort level immunopeptidomes to understand their individual presentation propensity. Thus, immunopeptidomics was used as circumstantial evidence of the biological availability of a mutated protein for presentation by HLA. First we mapped the mutational and immunopeptidome landscapes of a cohort of LUAD and LUSC patients. We then predicted HLA-restricted neoantigens using existing algorithms and used immunopeptidomic data from their individual tumours to filter those predictions on the basis of evidence that they could be presented. Neoantigen selection success was increased to 13%, with strong functional responses confirmed in 5 of 6 patients (83.3%) for our neoantigen panels selected for each donor. Finally, we used Bayesian modelling of the cohort’s immunopeptidomes to quantify how donor-specific presentation propensity and protein-level evidence combine to determine peptide presentation, providing a statistical framework for understanding our rules-based selection success.

These proof-of-concept results demonstrate how the information contained within the immunopeptidome has the potential to enhance proteogenomics strategies for identifying neoantigens for every patient according to their specific presentation propensity, enabling precision neoantigen selection that accounts for each patient’s unique presentation biology.

## Results

### A proteogenomics workflow for neoantigen identification

Our NSCLC cohort consisted of 24 patients, 15 LUAD (8 female, 7 male) and 9 LUSC (5 female, 4 male). Median age at diagnosis was 69 (See Table 1 and Supplementary Data S1). Tumour tissue and PBMCs were used for HLA typing, whole exome sequencing, RNA sequencing and mass spectrometry proteomics of the HLA immunopeptidome (Figure 1). To identify candidate neoantigens for each patient we developed a workflow that surveyed both the genomic and immunopeptidomic landscapes. Somatic missense variants called from the whole exome sequencing (WES, Supplementary Data S2 and S3) were used to generate a mutanome for each individual against which the HLA immunopeptidome could be searched for direct observation of neoantigens. Variants, gene expression and the patient HLA allotypes were also used for the prediction of putative neoantigens using existing tools^21^.

**Figure 1:**
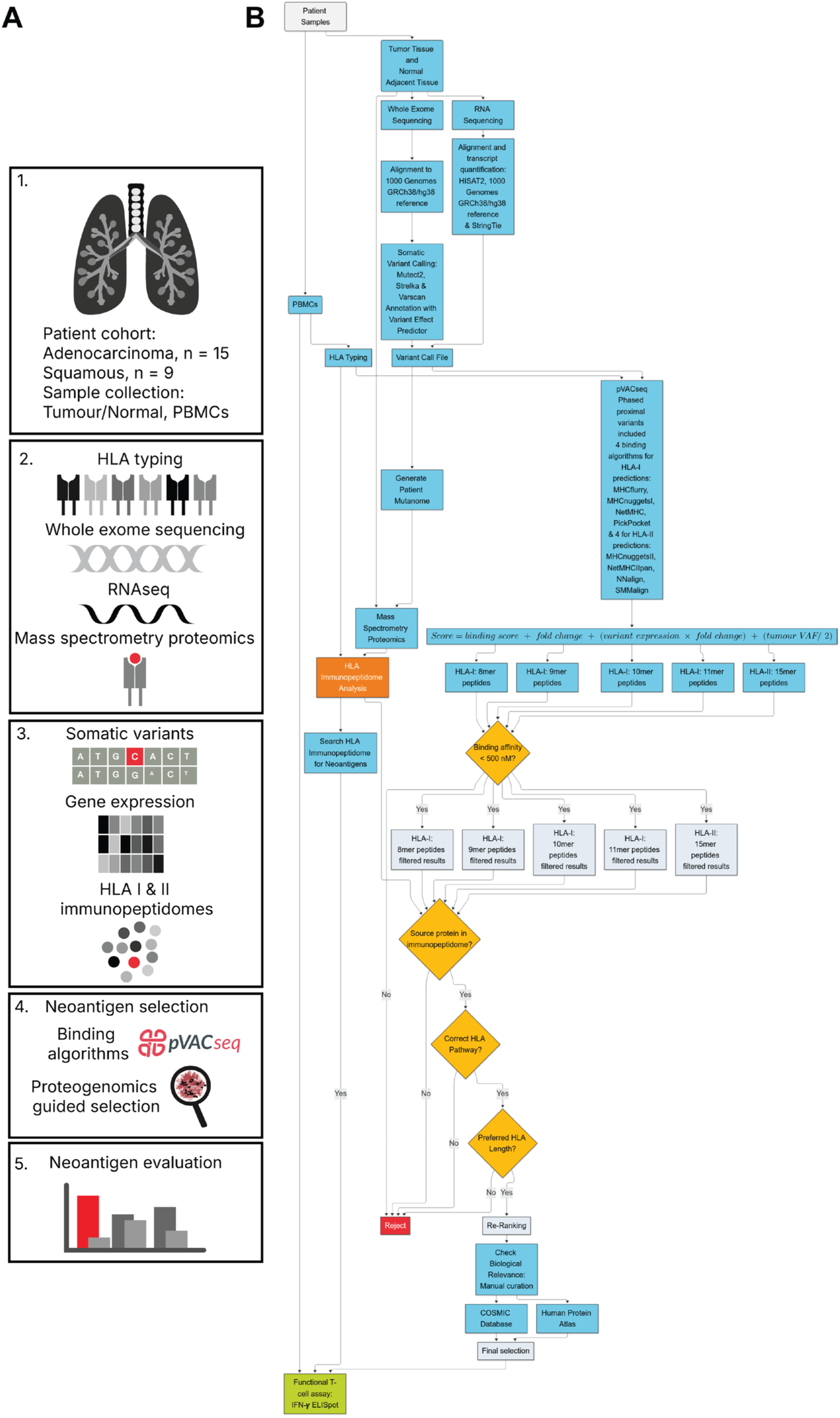
Integrated proteogenomics workflow. (A) HLA typing, whole exome sequencing, RNASeq and mass spectrometry-based proteomic of the HLA immunopeptidome were collected for 24 lung cancer patients providing mutational, gene expression and immunopeptidomic data from which to identify candidate neoantigens using binding algorithms and manual inspection of the combined proteogenomic data. (B) Flowchart of neoantigen selection workflow

**Table 1:** Clinical summary of patients in this study with non-small cell lung cancer.

| Donor | Smoking status | Cancer subtype |
| --- | --- | --- |
| A113 | Current smoker | Squamous |
| A114 | Ex smoker | Adenocarcinoma |
| A115 | Ex smoker | Squamous |
| A116 | Never smoker | Squamous |
| A117 | Current smoker | Adenocarcinoma |
| A118 | Ex smoker | Adenocarcinoma |
| A119 | Ex smoker | Adenocarcinoma |
| A120 | Ex smoker | Adenocarcinoma |
| A133 | Ex smoker | Squamous |
| A134 | Current smoker | Squamous |
| A136 | Never smoker | Adenocarcinoma |
| A137 | Ex smoker | Adenocarcinoma |
| A139 | Ex smoker | Adenocarcinoma |
| A140 | Ex smoker | Squamous |
| A141 | Ex smoker | Adenocarcinoma |
| A142 | Current smoker | Adenocarcinoma |
| A143 | Ex smoker | Adenocarcinoma |
| A144 | Ex smoker | Squamous |
| A145 | Ex smoker | Squamous |
| A146 | Current smoker | Adenocarcinoma |
| A147 | Current smoker | Adenocarcinoma |
| A148 | Ex smoker | Adenocarcinoma |
| A152 | Ex smoker | Squamous |
| A153 | Ex smoker | Adenocarcinoma |

### The mutational landscape of NSCLC in the studied cohort

To assess the likelihood of identifying HLA presented neoantigens we first examined the mutational landscape of the NSCLC cohort and found it to be consistent with previous reports^22,23^. Somatic variants were identified by WES of tumour and matched normal adjacent tissues. Tumour mutational burden (TMB) quantifies the number of mutations per million bases (Mb). From WES it is calculated as the number of variants divided by the size of the exome targets; here the target size was 35.7 Mb and the number of variants were either the total number of all variants, or only the protein coding missense variants: *N_vars_*/(35.7) = *N_vars_*/*Mb*.

This revealed that both cancer types have relatively high mutational burdens calculated from all variants, ranging from 27 to 280 mutations per Mb (Mt/Mb) with similar median mutational burdens of 109 Mt/Mb for LUAD and 104 Mt/Mb LUSC, but a broader range for LUAD (Figure 2 A, Supplementary Data S1). In terms of missense variants alone, this scales as ranging from 5 to 43 Mt/Mb and medians of 20 Mt/Mb for LUAD and 16 Mt/Mb LUSC

**Figure 2:**
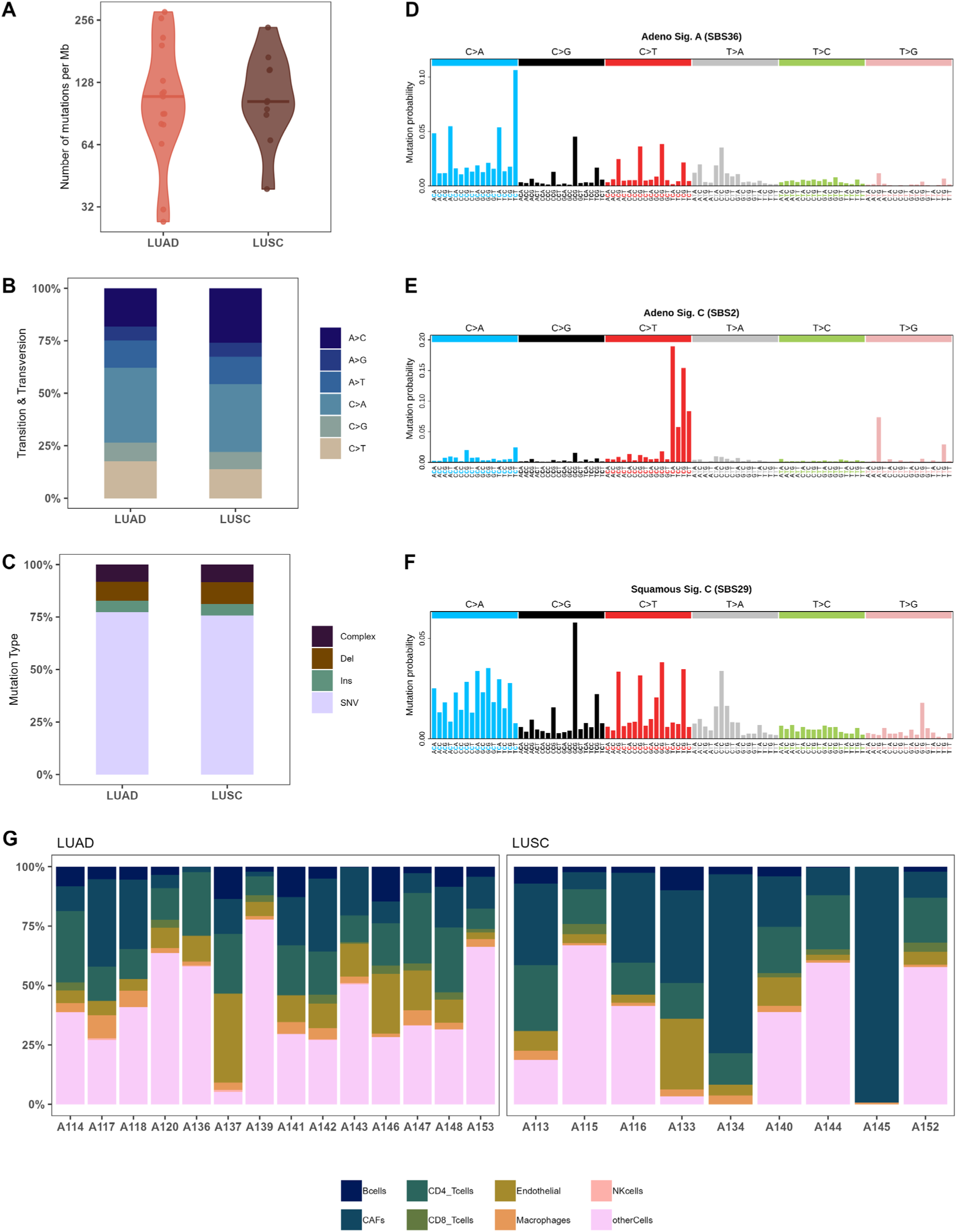
The mutational landscape of lung cancer in the studied cohort. (A) The mutational burden of each cancer type: squamous (n=9) and adenocarcinoma (n=15). (B) Mutation frequency of six transition and transversion categories for each cancer type. (C) Mutation frequencies each cancer type. (D-F) Mutational signatures identified in each cancer subtype. (G) The proportions of immune cells estimated from bulk tumor RNASeq in each tumour sample.

Approximately one third of all nucleotide transitions and transversions were C>A transversions in both LUSC and LUAD (Figure 2 B), a known mutational signature of smoking^22^. Of the 51,810 LUAD and 32,344 LUSC single nucleotide variants, approximately 20% were missense variants (10,565 LUAD, 6,772). These missense SNVs along with approximately 15% insertion/deletion variants (9,697 LUAD, 6,780 LUSC) predict amino acid changes at the protein level and are therefore potential sources of HLA neoantigens (Figure 2 C).

For each cancer subtype, patterns of single base substitutions created by the somatic mutations were extracted to identify mutational signatures that were fitted to those identified in COSMIC^24–26^ (Figure 2 D-F). LUAD signature A fit SBS36 indicating base excision repair deficiency characterised by C>A transversions. LUAD signature C fit SBS2, which is common in lung cancer and thought to indicate APOBEC cytidine deaminase activity as characterised by C>T transitions. LUSC signature C fit SBS29, another signature characterised by C>A transversions and linked to tobacco chewing.

In addition to examining the potential for neoantigen generation at the exon level, we sought to examine the potential for neoantigen recognition within the tumours using bulk gene expression data from RNAseq to assess the fractions of immune cells present in the tumours^27^ (Figure 2 G). This estimation also provides an indication of the tumour sample purity. All expressed genes not used as markers for immune cells are labelled as ‘otherCells’ and we would expect this category to comprise the largest proportion of cells in a tumour sample. Therefore if a sample has lower proportion of ‘otherCells’ it is indicative of a less pure tumour sample. For LUAD and LUSC, the median proportions of ‘otherCells’ are one third. In cases with very low proportions of ‘otherCells’ such as A134 and A145, the corresponding histology reports indicate these were fibrotic samples consistent with the very high proportions of cancer associated fibroblasts identified by RNAseq. However at the cohort level, proportions of T-cells estimated capable of responding to neoantigens presented by HLA were estimated with similar medians for CD4+ T-cells of 18% and 15% for LUAD and LUSC respectively, and medians for CD8+ T-cells of 2% and less than 1% for LUAD and LUSC respectively.

In summary, the mutational landscape of the NSLC cohort is characterised by a high tumour mutational burden in both cancer subtypes, the largest proportion of variants with the potential for generation of neoantigens arising from C>A transversions. Furthermore, gene expression data estimates the presence of limited populations of T-cells with the potential to recognise HLA presented neoantigens.

### The peptidome landscape of NSCLC in the studied cohort

Mass spectrometry proteomics of the HLA immunopeptidomes identified large distributions of peptides with their characteristic modes of 9 amino acids (AA) and 15 AA for class I and II HLA peptides respectively (Figure 3 A). Median class I immunopeptidome sizes were 5422 and 2998 for Adenocarcinoma and Squamous NSCLC respectively. Median class II immunopeptidome sizes were 2849 and 1125 for Adenocarcinoma and Squamous NSCLC respectively.

**Figure 3:**
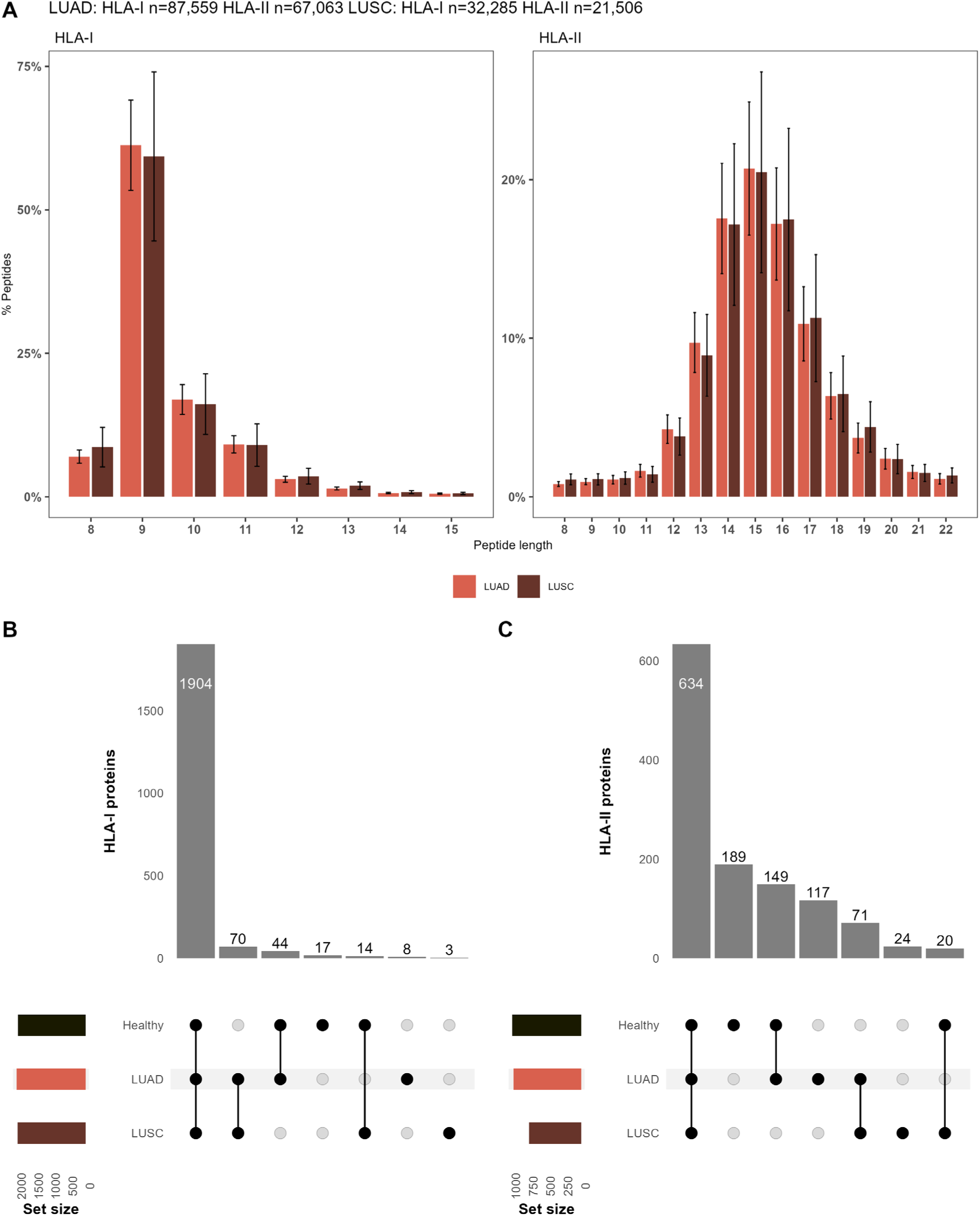
The peptidome landscape of lung cancer. (A) Length distributions of immunopeptides from tumour tissues. (B-C) Upset plots of proteins presented by HLA molecules (class I left, class II right) comparing proteins between cancer subtypes and healthy lung tissues from the HLA ligand atlas. Proteins were included if observed in at least two-thirds of the peptidomes for each cancer subtype.

### The lung cancer peptidome resembles the healthy lung tissue peptidome

We compared the distinct source protein populations yielding the class I and II HLA peptidomes between our LUAD, LUSC samples and healthy lung tissues from the Human HLA Ligand Atlas^10^ to examine their similarities and differences (Figure 3 B-C, Supplementary Data S5), considering only proteins present in at least two-thirds of our samples peptidomes. Our analysis suggests that healthy lung and tumour tissues immunopeptidomes sample largely the same protein populations. 92% of HLA-I proteins and 52% of HLA-II proteins were common to all three tissue types. The remaining proteins most likely represent experimental variation.

### Direct identification of a neoantigen in a LUAD patient immunopeptidome

Across the cohort of 24 patients we identified a single missense variant product by direct mass spectrometric observation in the class I HLA immunopeptidome of one LUAD patient (A147) (Figure S1-S3). This derived from a C>A variant in the ALYREF gene yielding an Asp10Tyr mutation in its protein product THO complex subunit 4 (Uniprot: Q86V81). This mutation yielded seven nested 15-18mer peptides with the mutation Y before the start of core sequence of MSLDDIIKL. No wild type peptides were observed for this protein in either the HLA I or II immunopeptidomes, suggesting this mutation altered either the binding affinity of these peptides or the source protein processing in the antigen processing pathway. The rarity of this observation is in keeping with estimates of frequencies in the order of 0.5% of missense variants encoding presented neoantigens^7,9^. The length of the ALYREF peptides suggested these may be class II HLA peptides that we had captured by chance in this assay. The motif most closely matched the patients HLA-DRB1*03:01 allotype with peptide AYKMDMSLDDIIKLN predicted as a weakly binding peptide^28^. We identified 1135 missense mutations for patient A147 (Supplementary Data S1). An estimate of 0.5% missense yielding neoantigens would represent 6 neoantigens. Here we observed one.

### A proteogenomics view of NSCLC in the studied cohort

Consistent with the view that non-silent mutations rarely encode HLA presented neoantigens^9^, we observed that LUAD and LUSC driver genes^29^ are not mutated and presented by HLA molecules with the same frequencies. Some drivers are frequently mutated, but rarely presented e.g. APC, whereas others are rarely mutated, but frequently present in the HLA immunopeptidomes e.g. KEAP1 (Figure 4 A). TP53 is both frequently mutated and presented in the class I HLA peptidomes of both NSCLC subtypes (Figure 4 A).

**Figure 4:**
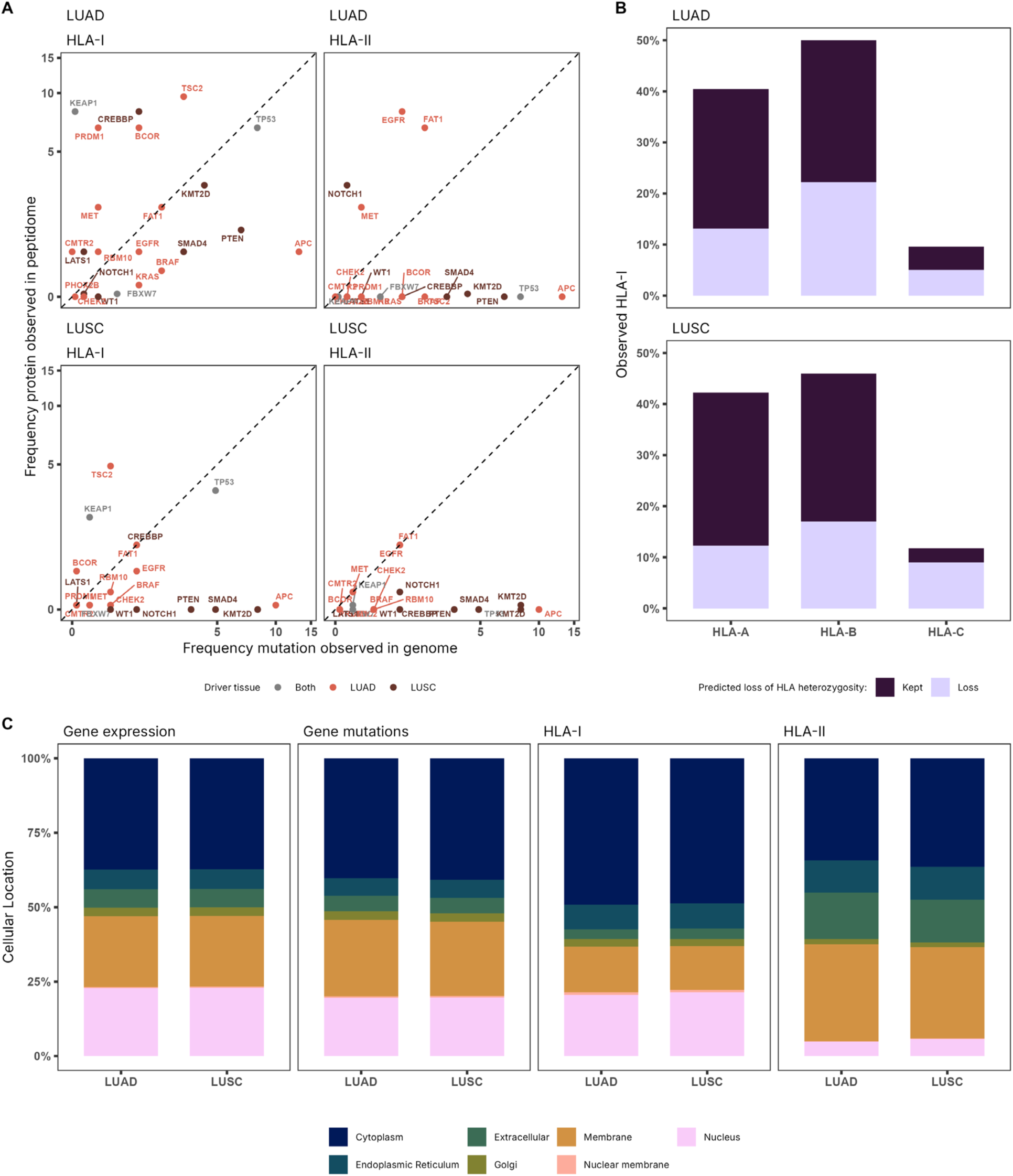
Integrating the mutational and immunopeptidome landscape reveals previously unclear relationships between mutations and peptide presentation. (A) Parity plots of the frequency of mutated initiating driver genes for each cancer type plotted against the frequency of observed presentation from the corresponding protein in either the HLA-I (left) or HLA-II peptidome (right). Parity of frequency is indicated by the dotted line. The colour indicates which cancer type the gene was identified as an initiating driver in^5^. (B) The relative proportions of immunopeptides assigned to each of the two HLA-A, B and C allotypes for heterozygous patients.^7,31,32^. The colour represents whether an allele is predicted to have loss of heterozygosity in the genome^30^. (C) Comparison of the frequency between the cellular compartments in which gene expression and somatic mutations occur, and those from which HLA peptides are observed in each cancer type.

We found that loss of class I HLA heterozygosity in the genome^30^ is reflected in the peptidome. In heterozygous patients, immunopeptides identified as presented by HLA molecules from the retained allele were observed at higher proportions in the peptidome than from the lost allele for HLA-A and B allotypes (Figure 4 B, Supplementary Data S4).

We also found that mutations are distributed across the cellular compartments at the same frequencies as the genes are expressed (Figure 4 C left), but the HLA pathways sample the compartments preferentially. Class I HLA immunopeptides are derived preferentially from nuclear and cytosolic proteins, whilst class II HLA immunopeptides are derived preferentially from membrane and extracellular proteins (Figure 4 C right).

Collectively, these observations imply that the likelihood of a putative neoantigen being presented by either HLA class is influenced by the cellular compartment origin of the source protein and secondly, putative neoantigens with motifs^31^ for the retained class I HLA allotypes are more likely to be presented than those from the lost allotype.

### Proteogenomics guided NSCLC neoantigen selection and testing

In selecting neoantigens we initially used the pVACseq tool to create a list of putative neoantigens for each patient and HLA allotype and different peptide lengths^33^. Briefly, we used pVACseq with the whole exome and transcriptome outputs and patient HLA allotypes to predict 8-11mer peptides for class I HLA and 15-mer peptides for class II HLA-DRB allotypes across eight binding algorithms. This combined genomic and binding score creates an overall score for each peptide (Details in Section 1.8.0.10). For our 24 patients this comprises 524 HLA class I tables and 74 HLA class II tables of ranked predictions. Discarding any prediction for a peptide with >500 nM binding affinity, pVACseq yielded 27,466 class I HLA and 127,015 class II HLA predicted neoantigen peptides (Supplementary Data S6-S8)

We were able to test predictions for six patients, but this still required selecting from thousands of possible candidate peptides. We consequently filtered the candidate peptides according to whether peptides arising from the gene product with a missense variant were already present in the patients’ respective class I or class II HLA peptidome (Supplementary Data S3) and according to HLA peptide length preferences^34^. This reduced the number of candidates to a few hundred peptides for each patient. We finally manually curated the ranked peptide candidates for biological relevance using auxiliary information from the literature, the Human Protein Atlas and COSMIC. (Figure 1, Section 1.8.0.10).

Our exploratory filtering process for candidate neoantigens can be summarised as: Does a missense mutation exist? Is there evidence that the mutated gene product enters the antigen processing pathway for presentation, and if so in which HLA pathway? Is the candidate neoantigen of the preferred HLA allotype length? Is the candidate neoantigen predicted to bind to the HLA allotype according to pVACseq? To preferentially select one neoantigen candidate over another in the final ranking we looked for evidence of biological relevance for the candidate in COSMIC^26^ and the HPA^35^.

As HLA peptidome observation took precedence over pVACseq rank, some candidates such as peptide 08-FAT1 were low ranking (70th percentile) but still with a predicted binding affinity lower than 500 nM (Table 2).

**Table 2:** Peptides yielding IFN-*γ* ELISPOT responses.

| Tissue | ID / HLA / Peptide Length<sup>a</sup> | Peptide<sup>b</sup> | Rank %<sup>c</sup> | Peptidome support<sup>d</sup> | Auxiliary support<sup>e</sup> | ELISpot response |
| --- | --- | --- | --- | --- | --- | --- |
| LUAD | A119 / DRB1*04:04 / 15 | 01-CANT1 | 11 | II | HPA | Strong |
| LUAD | A119 / HLA-A*31:01 / 10 | 12-PTPRT | 10 | I | COSMIC | Strong |
| LUAD | A147 / Observed / 15 | 01-ALYREF | - | I | - | Strong |
| LUAD | A147 / DRB1*04:01 / 15 | 08-FAT1 | 70 | I+II | COSMIC | Strong |
| LUAD | A147 / HLA-A*01:01 / 9 | 14-TP53 | 7 | I | COSMIC | Weak |
| LUAD | A148 / HLA-A*26:01 / 9 | 01-KMT2C | 20 | I | COSMIC | Strong |
| LUAD | A148 / DRB1*01:01 / 15 | 05-NT5E | 6 | I+II | HPA | Strong |
| LUSC | A134 / DRB1*01:03 / 15 | 06-KRT8 | 10 | I+II | - | Strong |
| LUSC | A144 / DRB1*04:01 / 15 | 04-FAT1 | 24 | I+II | COSMIC | Strong |
| LUSC | A144 / DRB1*04:01 / 15 | 08-NF1 | 3 | I+II | COSMIC | Strong |

<sup>a</sup>The patient ID, predicted HLA allotype for the peptide and peptide length. ALYREF was an observed peptide.
<sup>b</sup>The peptide identifier and gene name corresponding with those in Figure 5.
<sup>c</sup>Rank % is the rank of the peptide in the table for that Donor and HLA allotype, a lower rank corresponds with better pVACseq score as detailed in the materials and methods.
<sup>d</sup>Peptidome support indicates from which class HLA peptidome source protein peptides were observed.
<sup>e</sup>Auxiliary support indicates support for biological relevance either from the lung cancer associated proteins in the Human Protein Atlas (HPA) or the Top 20 mutated genes in COSMIC.

For six patients, 3 LUAD and 3 LUSC, we selected 9 to 14 putative neoantigens per patient (70 in total) and synthesised the specific putative HLA-I or HLA-II peptides in the mutant neoantigen and wildtype forms (Supplementary Data S9). Using their donor matched PBMCs, we identified nine strong neoantigen specific responses to putative neoantigens in five out of six patients, including for the directly observed ALYREF peptide (Figure 5 A-F, Table 2). This represents an 83% success in achieving a donor functional response and a 13% response rate in antigenic peptide selection, twice the genomics-based peptide prediction rate of 6% reported in the literature^15^. We observed responses to both class I and class II HLA candidate neoantigens in LUAD (Figure 5 A-C), but only class II HLA candidate neoantigens in LUSC (Figure 5 E-F).

**Figure 5:**
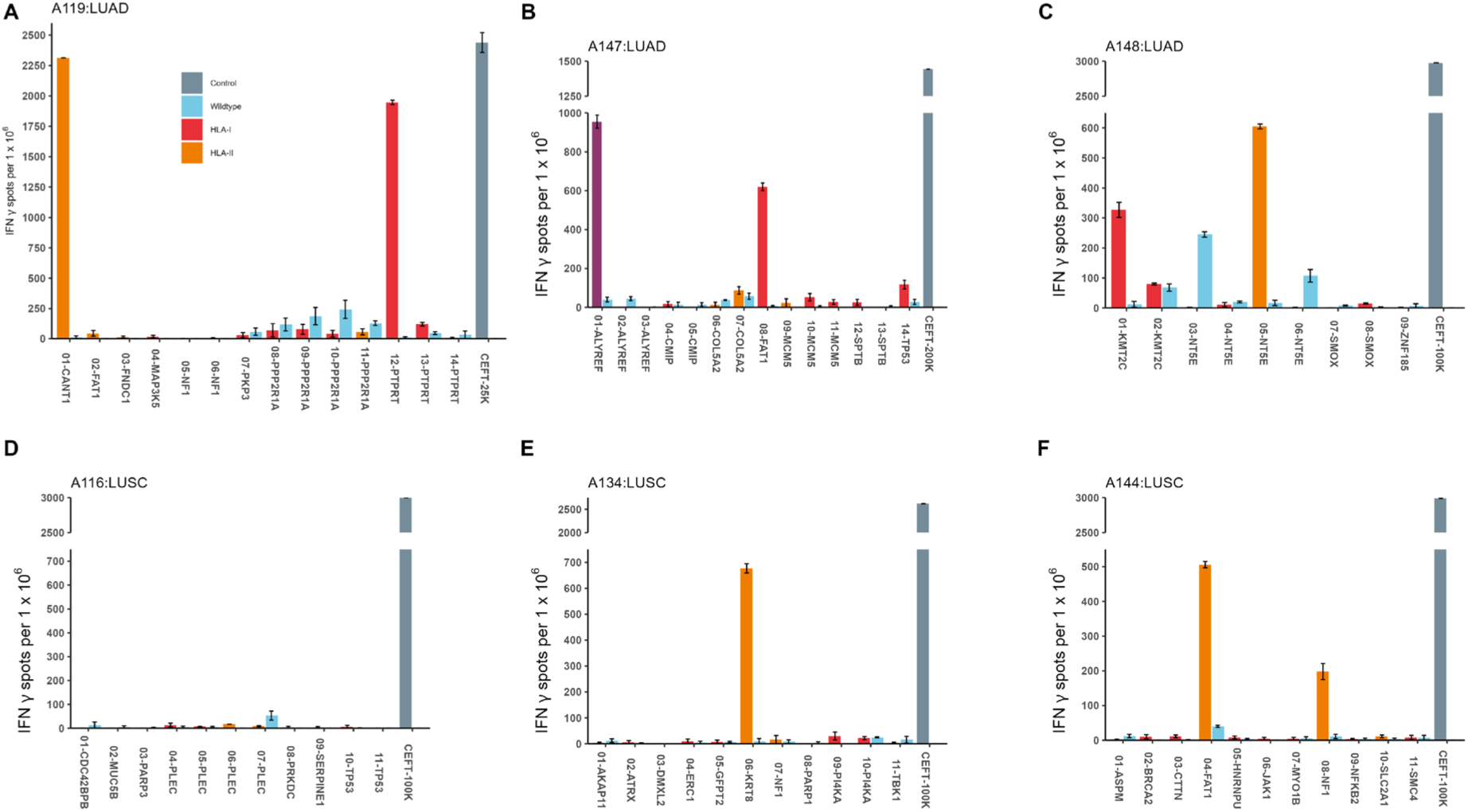
Proteogenomics guided NSCLC neoantigen selection identifies nine strong candidates (A-F) IFN-*γ* ELISPOT of putative neoantigens three LUAD and three LUSC patients donor matched PBMCs. Wildtype peptides are represented by blue bars, putative HLA-I neoantigens by red bars, putative HLA-II neoantigens by orange bars, the observed ALYREF neoantigen in purple and the CEFT control peptide mix in grey.

LUSC patient A116 yielded no responses (Figure 5 D).

### Model of peptide presentation

To retrospectively examine the mechanisms underlying our rules based selection approach and quantify the relative importance of factors affecting peptide presentation, we created Bayesian multilevel regression models of the NSCLC immunopeptidomes^36^. The goal was to understand the utility of immunopeptidomes in the context of neoantigen selection by quantifying what determines observed peptide presentation.

Several features have been identified as causal: protein abundance, protein hotspots, peptide length preferences of HLA molecules, peptide binding motifs and other features of antigen processing proteins such as tapasin, TAPBR and TAP on HLA-I peptide selection^37–41^.

Our findings indicated that driver genes do not correlate with peptide presentation (Figure 4 A) and that the presented proteome of LUAD, LUSC and healthy lung tissues are highly similar (Figure 3 B-C). Additionally, as others have also found, direct observations of neoantigens are rare^7,9^. Although not exclusively, we found that proteins were preferentially presented via HLA pathways corresponding with the cellular compartment they occupy (Figure 4 C).

To construct the models we began with the source proteins, which we did not observe directly. Proteins from the underlying proteome were latent variables from which peptides derived, bound to and were presented by HLA molecules. Proteins were represented in the form of peptide counts for each protein observed, for each HLA allotype and donor. We assumed that there were relationships between donor or NSCLC subtype cohort and protein properties e.g. abundance, with other peptides on binding to HLA and presentation, represented as a causal model by a directed acyclic graph (DAG) (Figure 6 A). Arrows indicated the direction of the causal effects the model assumed. The DAG indicated several sources of confounding, specifically the need for the model to adjust during fitting for protein, donor, cohort, peptide and HLA specific effects in estimating the probability of each peptide being presented. Within the limits of detection, a peptide was either presented or not, hence we assumed this outcome variable (yes or no) as being a Bernoulli distribution arising from a linear combination of our predictor variables: protein, donor, cohort, peptide and HLA properties. Importantly, having assumed the outcome distribution, Bayes theorem was used to pool observations across each subtype cohort and share information to adjust the estimation for each peptide as indicated by the DAG. Pooled adjustment helped to account for differing immunopeptidome sizes between donors and also enabled estimation of the probability of presentation of a peptide even if it was missing in any given immunopeptidome. These models quantified the relative importance of donor specific versus protein specific factors in determining peptide presentation.

**Figure 6:**
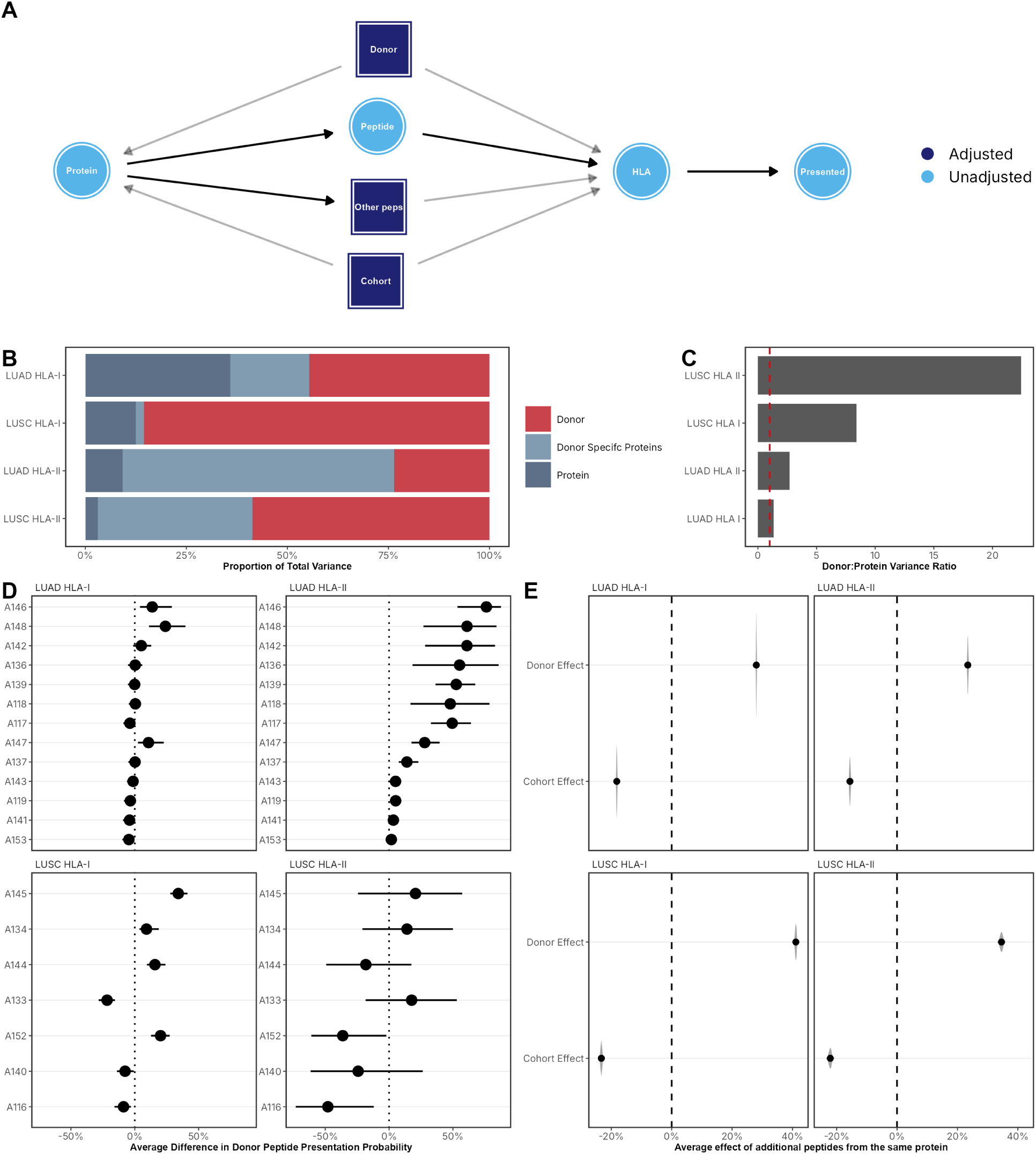
Bayesian multilevel regression model of HLA peptide presentation (A) Directed Acyclic Graph of HLA presentation for NSCLC cohorts. Variables adjusted for confounding in dark blue, unadjusted variables in light blue. (B) Contributions of donor differences, protein properties and donor specific proteins to the total variance in peptide presentation in Bayesian multilevel regression models of NSCLC immunopeptidomes. (C) Barplot of the ratio between donor and protein variance components in the peptide presentation models. The red dashed line indicates where these components are of equal value. (D) Comparison of the average propensity of donor to present peptides with respect to median presentation for each NSCLC subtype and HLA class. (E) Comparison of the average effect on the probability of presentation of an additional peptide from the same protein if a donor has already presented a peptide versus multiple peptides from that protein being presented across the cohort for each NSCLC subtype and HLA class.

Model diagnostics confirmed robust estimation with high signal-to-noise ratios (SNR > 10), adequate effective sample sizes, and good MCMC convergence across all models (Supplementary Tables S1-S2, Figures S4-S11).

To examine the relative importance of donor differences, protein properties and donor-specific protein preferences, we calculated the contribution to the total variance of each model component (Figure 6 B). We found moderately greater contributions of donor effects to protein effects for LUAD HLA-I and HLA-II presentation, ratios of 1.3 and 3 times greater respectively (Figure 6 C). For the LUSC cohort, donor effects were much larger contributors to presentation variance than protein effects for both HLA-I and II, ratios of 8 and 22 times respectively (Figure 6 C). Donor-specific protein contributions varied and were greatest for HLA-II immunopeptidomes, and was the largest variance component for LUAD HLA-II.

To further quantify these effects, we calculated the average difference in peptide presentation probability for each donor relative to median donor presentation (Figure 6 D). For LUAD, most donors showed HLA-I presentation close to the median, but differences in the order of 50% for some donors with respect to median presentation for HLA-II, albeit with much greater uncertainty. The model estimated donors such as A148 and A147 were more likely to present peptides, despite A148 and A147 having only the 6th and 10th highest number of HLA-I and 8th and 9th number of HLA-II peptide observations respectively (Supplementary Data S1). The estimates for LUSC donors showed greater variation and uncertainty, with most donors below the median. Donor A116 had the greatest number of LUSC HLA-I peptide observations, and 3rd greatest LUSC HLA-II peptide observations (Supplementary Data S1), but the model estimated A116 below the median for presentation for both HLA classes in LUSC, and we were unable to select a reactive neoantigen peptide for this donor.

In our rules-based selection of neoantigens we ranked a prediction more highly if a donor presented peptides from the same protein as the putative neoantigen (Figure 1 B). To quantify this effect and examine our rules-based assumption, we calculated the average effect on peptide presentation of observing multiple peptides from the same protein across each NSCLC subtype cohort versus the effect of observing more than one peptide from the same protein presented by a donor.

The model identified competing effects such that when a protein presented multiple peptides across the cohort, this decreased the probability of observing an additional peptide by between 18-23% depending on NSCLC subtype and HLA class (Figure 6 E). In contrast, a donor presenting one peptide from a protein increased the probability of observing another peptide from the same protein by between 23-41% depending on NSCLC subtype and HLA class (Figure 6 E), supporting our rules-based assumption. The model quantified that donor-specific protein preferences substantially influenced presentation probability, but also revealed that cohort-level presentation patterns created competing effects.

In summary, by pooling information across immunopeptidomes in the NSCLC cohorts, the models identified NSCLC subtype variation and donor-specific effects on peptide presentation: each donor’s general propensity to present peptides and their protein-specific preferences.

## Discussion

The strategy of using HLA presented peptides as a basis for immunotherapy is long standing^42,43^. Researchers have sought to identify either peptides common to a cancer type, so-called tumour associated antigens, or peptides unique to a patient’s tumour, so called neoantigens. Here we sought to improve our ability to identify HLA presented neoantigens in two NSCLC sub-types using a proteogenomics approach that combines exome sequencing, transcriptomics, mass spectrometry immunopeptidomics and mechanistic modelling. In tumour samples from cohort of 24 NSCLC patients we found relatively high mutational burdens with exonic mutations characterised by a predominance of C>A transversions and containing small populations of T-cells. Consistent with previous reports^8,9^ using mass spectrometry immunopeptidomics, we only directly identified one neoantigen amongst tens of thousands of peptide identifications. However, following a rules based approach, we utilised the remaining observations to inform our selection of neoantigens from ranked lists generated by in silico prediction algorithms^33^ to the extent that we were able to identify positive functional assay neoantigens for 5 out of the 6 patients (83%) we were able to test. This included a positive response for the directly observed neoantigen. These findings also represent a two-fold improvement over previous reports for neoantigen prediction and identification^15,16^. We then applied Bayeisan inference to retrospectively examine our rules based neoantigen selection and quantify patient specific differences in peptide presentation propensity.

Although identifying patient specific neoantigens and was the primary goal of our study, the data included some interesting related observations: With the exception of TP53 and FAT1 in the HLA-I and HLA-II immunopeptidomes respectively, there was no correlation between driver gene mutation frequency and their peptide presentation frequency. This provides some circumstantial support for these two genes as sources for neoantigens^44,45^. TP53 mutations can be either truncal or late-stage^46^, but a third of TP53 mutations occur in so-called hotspot regions^47^, making them of interest both for early detection and as targets for immunotherapy^48,49^. Overall we found that the tumour peptidomes contained peptides derived from the same source proteins as healthy tissue. Furthermore, although somatic mutations are not random, as seen in the mutational signatures and driver genes, their distribution amongst cell compartments corresponds with gene expression frequency. There is no enrichment for mutations in genes expressing proteins in specific cell compartments. This implies a connection between the cell compartment from where the source protein derives and the subsequent HLA antigen processing pathway it primarily feeds, consistent with previous reports^50^. TP53 is a predominantly a nuclear protein, whilst FAT1 is predominantly extracellular, hence their higher frequencies in HLA I and II immunopeptidomes respectively. Whilst this might seem tautological, it does indicate that it would be unwise to preferentially select class I neoantigen predictions for FAT1 and vice versa for TP53, and yet this is not explicitly considered in existing neoantigen prediction algorithms. Hence, we selected source protein cell compartment as a relevant neoantigen parameter in our rules based approach. This choice was supported by the mechanistic model prediction of an increased probability of observing another peptide from the same protein if a patient has presented one peptide already by between 23-41%. However, our mechanistic model also identified competing effects between proteins that were widely presented in NSCLC subtype and those proteins specific to each donor, highlighting the need for patient specific predictions that aren’t solely reliant on general HLA peptide characteristics.

Loss of class I HLA heterozygosity in the genome was reflected in the proportions of peptides observed for each HLA-I allotype, and although we did not use this information to select neoantigens, this might be another useful parameter when ranking candidates.

There are various limitations in our study which might be addressed by future studies. Perhaps most significantly from a methodological perspective, mass spectrometry as a technique does not have the amplification step found in many genomic sequencing methodologies. Therefore the strength of the input signal arises almost entirely from sample quality and preparation and sensitivity is determined by the mass spectrometer itself. The complexity of the input mixture and the differential ability of peptides to ionise, along with their relative abundances all affect what fraction of the immunopeptidome is observed. Various single molecule technologies are being developed that may address this problem, of which pore-based technologies, possibly in combination with fluorescence fingerprinting, seem well suited to identification of short peptides^51–53^. Sequencing peptides using pore technologies offers the tantalising prospect of providing much greater coverage of the immunopeptidome, and therefore direct observation of neoantigens. There are many challenges to this approach, not least post-translational modifications and the non-polar nature of protein peptides, but much progress has already been made^54–56^.

In our study we only considered canonical neoantigens arising from missense variants. This was a limitation largely arising from choosing whole exome sequencing, but there is increasing evidence for non-canonical antigens arising from non-coding regions of the genome^57–59^. There are several sources of non-canonical peptides such as circRNA translation, unannotated open reading frames (nuORFs), lncRNA and alternative reading frames. As yet, clinical efficacy is unknown and it is also unclear how abundant non-canonical peptides are in the immunopeptidome due their precursor instability and/or relative abundance^60^. Likewise, detection of non-canonical peptides is further hampered by the increase in search space they represent and subsequent impact on detection sensitivity, though this is being addressed^61^. Immunopeptidomic identification of circRNA and cancer specific nuORFs peptides have been reported^63^ and immune responses in mice to pancreatic cancer nuORF peptides^64^. Therefore we’d expect inclusion of non-canonical targets to be part of future immunopeptidomic studies.

Here we have identified potential candidates for personalised vaccines that elicit strong positive responses in functional T-cell assays, however further evaluation is required to determine their clinical efficacy, and their effectiveness as vaccines cannot be guaranteed. The stage of the cancer at which the patient receives the vaccine may be crucial for efficacy. Chronic neoantigen exposure driving T-cells to dysfunctional states, late-differentiated T-cells dominating the tumour microenvironment, and loss of HLA heterozygosity are all reasons NSCLC may become harder to treat with neoantigen vaccines at later stages^6,65–67^. Heterogeneity in NSCLC tumours is likely to influence the efficacy of neoantigen based vaccines^29^.

Personalised neoantigen vaccines are already being trialled for the treatment of melanoma, glioblastoma and pancreatic cancer^68–70^. These trials rely on the delivery of mRNA containing a number of long sequences predicted to be processed into the final HLA presented neoantigens. The vaccine response rate is in currently the order of 50% of patients when multivalent epitope vaccines in combination with an checkpoint inhibitor adjuvant are used^71^. So whilst these results are extremely promising, there is clearly room for improvement, including in the neoantigen selection process. Immunogenic peptides are predicted by algorithms that incorporate machine learnt parameters such as peptide binding affinity^72^ or proteasomal cleavage^73^, or more recently using machine learning to identify features such as protein hotspots from large mass spectrometry immunopeptidomics datasets^17,18^.

The principal difference in our approach is one of tactics rather than strategy. Our tactical difference being to look at which proteins yield peptides presented by HLA molecules and then manually identifying supporting evidence for each neoantigen candidate protein in the literature. To understand the mechanistic basis for this approach, we retrospectively applied Bayesian inference to quantify how donor-specific and protein-specific factors contribute to peptide presentation. This analysis validated our empirical observation that donor-presented source proteins are informative for candidate ranking, while revealing competing cohort-level effects that could inform parameter weighting in future automated selection algorithms. This tactic has some similarity to the ‘Tübingen approach’ for identification of tumour associated neoantigens which uses mass spectrometry proteomics identifications of HLA peptides to rank candidates^74^, as used in the glioblastoma vaccine^69^.

Related immunopeptidomic-guided approaches have been reported. Tokita et al. used pooled wildtype immunopeptidomes as “surrogate” immunopeptidomes to filter neoantigen candidates against exonic mutations, successfully identifying two neoantigens in colorectal cancer^75^. A large-scale immunopeptide-guided selection study used machine learning to analyse approximately 0.3 million peptides^19^. In concordance with estimates that only 0.5% of mutations yield presented neoantigen peptides^9^, they predicted 0.7% of peptides as immunogenic (2,523 peptides total: 1,476 canonical and 1,047 noncanonical), with three canonical peptides validated. Our approach differs in using patient-specific tumour immunopeptidomes rather than surrogate immunopeptidomes, systematically testing multiple candidates per patient (70 peptides across 6 patients) and using mechanistic modelling to identify relevant parameters for machine learning models.

Although still far from perfect with 87% of our predictions failing, our approach outperformed current machine learning models and, in the tested patients, identified peptides that elicited a strong functional response in ∼83% of cases tested. Perhaps most importantly, our findings indicate that patient specific knowledge beyond their mutational differences, incorporating knowledge of each individual’s HLA-presented peptides are critical parameters in improving neoantigen selection workflows and precision therapies.

## Materials and Methods

### Ethics statement

This study was performed in accordance with the Declaration of Helsinki. Ethical approval was obtained from the local research ethics committee (LREC reference 14-SC-0186 150975) and written informed consent was provided by the patients.

### Tissue preparation

Tumours were excised from resected lung tissue post-operatively by pathologists and processed either for histological evaluation of tumour type and stage, or snap frozen at −80°C. Whole blood samples were obtained, and PBMCs were isolated by density gradient centrifugation over Lymphoprep prior to storage at −80°C.

### HLA typing

HLA typing was performed by Next Generation Sequencing by the NHS Blood and Transplant Histocompatibility and Immunogenetics Laboratory, Colindale, UK.

### DNA and RNA extraction

DNA and RNA were extracted from tumor tissue that had been obtained fresh and immediately snap frozen in liquid nitrogen. Ten to twenty 10 µm cryosections were used for nucleic acid extraction using the automated Maxwell® RSC instrument (Promega) with the appropriate sample kit and according to the manufacturer’s instructions: Maxwell RSC Tissue DNA tissue kit and Maxwell RSC simplyRNA tissue kit, respectively. Similarly, DNA was extracted from snap frozen normal adjacent tissue as described above. DNA and RNA were quantified using Qubit fluorometric quantitation assay (ThermoFisher Scientific) according to the manufacturer’s instructions. RNA quality was assessed using the Agilent 2100 Bioanalyzer generating an RNA integrity number (RIN; Agilent Technologies UK Ltd.).

### Whole exome sequencing

The tumor and normal adjacent samples were prepared using SureSelect Human All Exon V7 library (Agilent, Santa Clara USA). 100 bp paired end reads sequencing was performed using the Illumina NovaSeq 6000 system by Edinburgh Genomics (Edinburgh, UK) providing ∼100X depth. Reads were aligned to the 1000 genomes project version of the human genome reference sequence (GRCh38/hg38) using the Burrows-Wheeler Aligner (BWA; version 0.7.17) using the default parameters with the addition of using soft clipping for supplementary alignments. Following GATK Best Practices, aligned reads were merged^76^, queryname sorted, de-duplicated and position sorted^77^ prior to base quality score recalibration^78^.

### Tumour purity and ploidy

Tumour purity and ploidy were estimated from WES using the ASCAT R package v3.3.0 following the recommended steps on the Github repository: https://github.com/VanLoo-lab/ascat^79^.

### Somatic variant calling

Somatic variant calling was performed using three variant callers: Mutect2 (version 4.1.2.0)^80^, Varscan (version 2.4.3)^81^, and Strelka (version 2.9.2)^82^. For Mutect2, a panel of normals was created using 40 samples (20 male and 20 female) from the GBR dataset. Variants were combined using gatk GenomeAnalysisTK (version 3.8-1) with a priority order of Mutect2, Varscan, Strelka. Variants were then left aligned and trimmed, and multi-allelic variants split^83^. Hard filtering of variants was performed such that only variants that had a variant allele fraction > 5%, a total coverage > 20 and variant allele coverage > 5 were kept. Filtered variants were annotated using VEP (version 97)^84^ and with their read counts (https://github.com/genome/bam-readcount) to generate the final filtered and annotated variant call files (VCF).

### RNA sequencing

Samples were prepared as TruSeq stranded mRNA libraries (Illumina, San Diego, USA) and 100bp paired end sequencing was performed using the Illumina NovaSeq 6000 system by Edinburgh Genomics (Edinburgh, UK). Raw reads were pre-processed to using fastp (version 0.20.0)^85^. Filtered reads were aligned to the 1000 genomes project version of the human genome reference sequence (GRCh38/hg38) using hisat2 (version 2.1.0)^86^, merged and then transcripts assembled and gene expression estimated with stringtie2 (version 1.3.5)^87^ using reference guided assembly.

### Mutanome generation

The annotated and filtered VCFs were processed using Variant Effect Predictor (version 97)^84^ plugin ProteinSeqs to derive the amino acid sequences arising from missense mutations for each sample for use in immunopeptide analyses.

### Neoantigen prediction

Variant call files were prepared for the pvacseq neoantigen prediction pipeline (version 1.5.10)^21,33^ by adding tumor and normal DNA coverage, and tumor transcript and gene expression estimates using vatools (version 4.1.0) (http://www.vatools.org/). Variant call files of phased proximal variants were also created for use with the pipeline^88^. Prediction of neoantigens arising from somatic variants was then performed using pvacseq with the patient HLA allotypes to predict 8-11mer peptides for class I HLA and 15-mer peptides for class II HLA-DRB allotypes. Four binding algorithms were used for class I predictions (MHCflurry, MHCnuggetsI, NetMHC, PickPocket) and four for class II predictions (MHCnuggetsII, NetMHCIIpan, NNalign, SMMalign). Unfiltered outputs were post-processed in R^89^ and split into individual tables for each peptide length and HLA allotype for each patient, and each table was then ranked according to the pvacseq score, where:

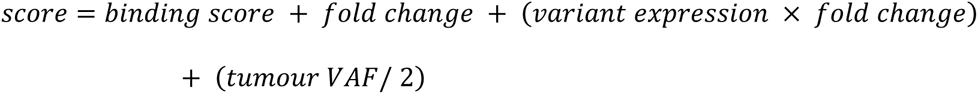

Here *binding score* is 1/median neoantigen binding affinity, *fold change* is the difference in median binding affinity between neoantigen and wildtype peptide (agretopicity).

Each table was then filtered according to whether wildtype peptide(s) from the same protein as predicted neoantigen was present in the individual’s peptidome, and further filtered manually according to biological relevance e.g. the ontology of the protein and its likely presence in the relevant HLA pathway, for example a cytoplasmic resident protein would be considered more likely to yield a HLA-I neoantigen than a HLA-II one. The Human Protein Atlas list of 354 genes identified for unfavourable prognosis in lung cancer, the COSMIC top 20 mutated genes and literature searches were also used as a screen for genes/proteins/peptides of biological relevance.

### Immunopeptidomics

Snap frozen tissue samples were briefly thawed and weighed prior to 30s of mechanical homogenization (Fisher, using disposable probes) in 4 mL lysis buffer (0.02M Tris, 0.5% (w/v) IGEPAL, 0.25% (w/v) sodium deoxycholate, 0.15mM NaCl, 1mM EDTA, 0.2mM iodoacetamide supplemented with EDTA-free protease inhibitor mix). Homogenates were clarified for 10 min at 2,000g, 4°C and then for a further 60 min at 13,500g, 4°C. 2 mg of anti-MHC-I mouse monoclonal antibodies (W6/32) covalently conjugated to Protein A sepharose (Repligen) using DMP as previously described^90,91^ were added to the clarified supernatants and incubated with constant agitation for 2 h at 4°C. The captured MHC-I/*β*_2_m/immunopeptide complex on the beads was washed sequentially with 10 column volumes of low (isotonic, 0.15M NaCl) and high (hypertonic, 0.4M NaCl) TBS washes prior to elution in 10% acetic acid and dried under vacuum. The MHC-I-depleted lysate was then incubated with anti-MHC-II mouse monoclonal antibodies (IVA12) and MHC-II bound peptides were captured and eluted in the same conditions.

Immunopeptides were separated from MHC-I/*β*_2_m or MHC-II heavy chain using offline HPLC on a C18 reverse phase column, as previously described^90^. Briefly, dried immunoprecipitates were reconstituted in buffer (1% acetonitrile,0.1% TFA) and applied to a 10cm RP-18e 100-4.6 chromolith column (Merck) using an Ultimate 3000 HPLC equipped with UV monitor. Immunopeptides were then eluted using a 15 min 0-40% linear acetonitrile gradient at a flow rate of 1 mL/min. Peptide fractions were eluted and pooled at between 0 and 30% acetonitrile, and the *β*_2_m and MHC heavy chains eluted at >40% acetonitrile.

HLA peptides were separated by an Ultimate 3000 RSLC nano system (Thermo Scientific) using a PepMap C18 EASY-Spray LC column, 2 µm particle size, 75 µm x 75 cm column (Thermo Scientific) in buffer A (0.1% Formic acid) and coupled on-line to an Orbitrap Fusion Tribrid Mass Spectrometer (Thermo Fisher Scientific,UK) with a nano-electrospray ion source. Peptides were eluted with a linear gradient of 3%-30% buffer B (Acetonitrile and 0.1% Formic acid) at a flow rate of 300 nL/min over 110 minutes. Full scans were acquired in the Orbitrap analyser using the Top Speed data dependent mode, performing a MS scan every 3 second cycle, followed by higher energy collision-induced dissociation (HCD) MS/MS scans. MS spectra were acquired at resolution of 120,000 at 300 m/z, RF lens 60% and an automatic gain control (AGC) ion target value of 4.0e5 for a maximum of 100 ms. MS/MS resolution was 30,000 at 100 m/z. Higher energy collisional dissociation (HCD) fragmentation was induced at an energy setting of 28 for peptides with a charge state of 2–4, while singly charged peptides were fragmented at an energy setting of 32 at lower priority. Fragments were analysed in the Orbitrap at 30,000 resolution. Fragmented m/z values were dynamically excluded for 30 seconds. Data desposited in PRIDE^92^, see Data Availability for details.

### Proteomic data analysis

Raw spectrum files were analyzed using Peaks Studio 10.0 build 20190129^93,94^ and the data processed to generate reduced charge state and deisotoped precursor and associated product ion peak lists which were searched against the UniProt database (20,350 entries, 2020-04-07) plus the corresponding mutanome for each sample (∼1,000-5,000 sequences) and contaminants list in unspecific digest mode. Parent mass error tolerance was set a 5ppm and fragment mass error tolerance at 0.03 Da. Variable modifications were set for N-term acetylation (42.01 Da), methionine oxidation (15.99 Da), carboxyamidomethylation (57.02 Da) of cysteine. As previously described, carbamidomethylated cysteines were treated as variable modifications due to the low concentration of 0.2 mM of iodoacetamide used in the lysis buffer to inhibit cysteine proteases^95^. A maximum of three variable modifications per peptide was set. The false discovery rate (FDR) was estimated with decoy-fusion database searches^93^ and were filtered to 1% FDR. Downstream analysis and data visualizations of the Peaks Studio identifications was performed in R using associated packages^89,96^.

### Immunopeptide HLA assignment

Identified immunopeptides were assigned to their HLA allotype for each patient using motif deconvolution tools and manual inspection. For class I HLA peptides initial assignment used MixMHCp (version 2.1)^7,31^ and for class II HLA peptides initial assignment used MoDec (version 1.1)^32^. Downstream analysis and data visualizations was performed in R using associated packages^89,96,97^.

### Synthetic peptides

Peptides for functional T-cell assays and spectra validation were synthesised using standard solid phase Fmoc chemistry (Peptide Protein Research Ltd, Fareham, UK).

### Functional T-cell assay

PBMC (2×10^6^ per well) were stimulated in 24-well plates with peptide (individual/pool) plus recombinant IL-2 (R&D Systems Europe Ltd.) at a final concentration of 5µg/mL and 20IU/mL, respectively, and incubated at 37°C with 5% CO2; final volume was 2mL. Media containing additional IL-2 (20IU/mL) was refreshed on days 4, 6, 8 and 11 and on day 13 cells were harvested. Expanded cells (1×10^5^ cell/well) were incubated in triplicate with peptide (individual) at 5µg/mL final concentration for 22 hours at 37°C in 5% CO2; phytohemagglutinin (PHA; Sigma-Aldrich Company Ltd.) and CEFT peptide mix (JPT Peptide Technologies GmbH, Berlin, Germany), a pool of 27 peptides selected from defined HLA Class I- and II-restricted T-cell epitopes, were used as positive controls. Spot forming cells (SFC) were counted using the AID ELISpot plate reader system ELR04 and software (AID Autoimmun Diagnostika GmbH) and positivity calling for ELISpot data used the runDFR(x2) online tool (http://www.scharp.org/zoe/runDFR/). Downstream analysis and data visualizations was performed in R using associated packages^89,96^.

### Peptide presentation model

Peptide presentation was modelled in R using Bayesian Multilevel Regression Models^36^. For all simulations the Monte Carlo Markov Chain settings were:

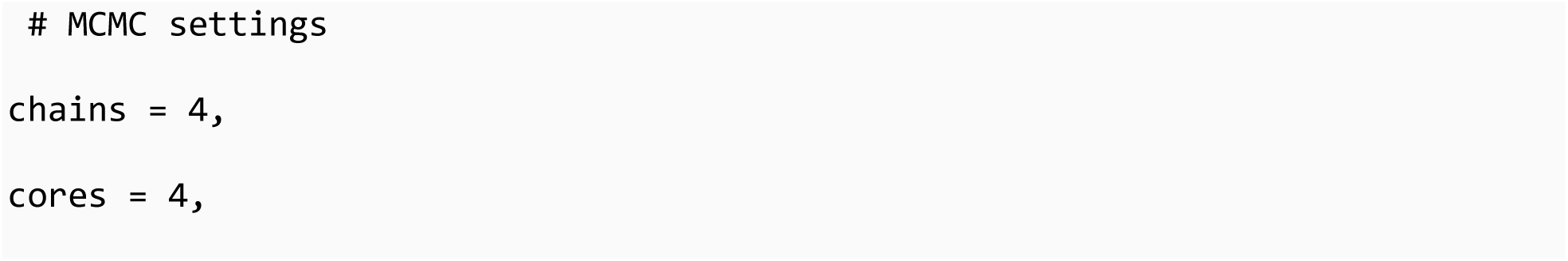
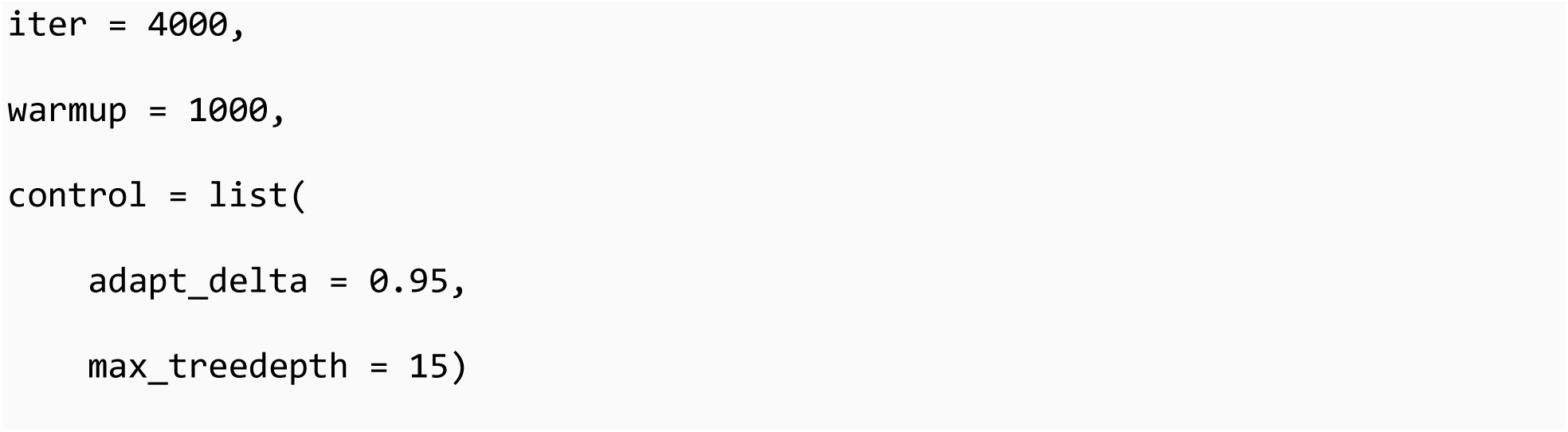

Model quality control was assessed using trace plots to assess convergence and effective parameter sampling (Figure S4-S7), and caterpillar plots to assess the uncertainty of model parameters (Figure S8-S11).

### HLA Class I model

A peptide is either presented or not, so peptide presentation is 1 or 0. The model is therefore defined using a Binomial distribution with a logistic link function:

~~~
peptide_presentation ∼ Binomial(1, p) logit(p) = η
~~~

The linear predictor η is defined as:

~~~
η = β₀ + β₁·binding_pre_threshold + β₂·binding_post_threshold +
β₃·length_diff_from_9 + Σⱼ β₁ⱼ·(binding_pre_threshold × I(hla = j)) + Σⱼ
β₂ⱼ·(binding_post_threshold × I(hla = j)) + Σⱼ β₃ⱼ·(length_diff_from_9 ×
I(hla = j)) + β₇·z_sqrt_peptides_donor + β₈·z_sqrt_peptides_cohort + u_donor
+ u_donor:uniprot_id + u_uniprot_id
~~~

Where:

- `j` is the index for each HLA allotypes (`j = 1, 2, …, J`)
- `I(hla = j)` equals 1 when the observed peptide has HLA allotype `j`, 0 otherwise
- `β₁ⱼ, β₂ⱼ, β₃ⱼ` are HLA-specific interaction coefficients

This allows binding and length preferences to vary by HLA allotype.

There are three levels of random effects:

1. Donor-level effects: `u_donor ∼ Normal(0, σ²_donor)`
2. Donor-protein interaction effects: `u_donor:protein ∼ Normal(0, σ²_donor:protein)`
3. Protein-level effects: `u_protein ∼ Normal(0, σ²_protein)`

Partial pooling via these random effects helps to account for donor heterogeneity and any protein-specific effects.

The variables are:

HLA-I binding scores were estimated using MixMHCpred3.0 and the resulting `score_best_allele` scores for each peptide were centred and split around 0 to capture strong and weak binding effects.

- `binding_pre_threshold`: Weak binding peptide score
- `binding_post_threshold`: Strong binding peptide score
- `length_diff_from_9`: Difference in peptide length from 9 amino acids
- `hla`: HLA allotype
- `z_sqrt_peptides_donor`: Standardised square root of number of peptides per protein observed per donor
- `z_sqrt_peptides_cohort`: Standardised square root of number peptides peptides per protein observed per cohort
- `donor`: Donor identifier
- `uniprot_id`: UNIPROT protein identifier

Prior Distributions:

- `β`₀ `∼ Normal(-1, 1)` Intercept
- `β₁ ∼ Normal(1, 0.5)` binding_pre_threshold (main effect)
- `β₂ ∼ Normal(1, 0.5)` binding_post_threshold (main effect)
- `β₃ ∼ Normal(0, 0.5)` length_diff_from_9 (main effect)
- `β₁ⱼ ∼ Normal(0, 0.5)` binding_pre_threshold:hla interactions
- `β₂ⱼ ∼ Normal(0, 0.5)` binding_post_threshold:hla interactions
- `β₃ⱼ ∼ Normal(0, 0.5)` length_diff_from_9:hla interactions
- `β`₇ `∼ Normal(0.5, 0.8)` z_sqrt_peptides_donor
- `β`₈ `∼ Normal(-0.6, 0.8)` z_sqrt_peptides_cohort
- `σ_donor ∼ Exponential(2)` Donor random effect SD
- `σ_donor:uniprot_id ∼ Exponential(2)` Donor-protein random effect SD
- `σ_uniprot_id ∼ Exponential(2)` Protein random effect SD

### HLA Class II model

As for HLA-I, peptide is either presented or not, so peptide presentation is 1 or 0. The model is therefore defined using a Binomial distribution with a logistic link function:

~~~
peptide_presentation ∼ Binomial(1, p) logit(p) = η
~~~

The linear predictor η is defined as:

~~~
η = β₀ + Σⱼ sⱼ(binding_score_centered) × I(hla = j) +
β₁·length_diff_from_15 + Σⱼ β₁ⱼ·(length_diff_from_15 × I(hla = j)) +
β₂·z_sqrt_peptides_donor + β₃·z_sqrt_peptides_cohort + u_donor +
u_donor:uniprot_id + u_uniprot_id
~~~

For HLA-II peptides, MixMHC2pred was used and produces scores as a binding rank and these were modelled as smooth splines:

~~~
Σⱼ sⱼ(binding_score_centered) × I(hla = j)
~~~

Where:

- `sⱼ(·)` are HLA allotype specific thin plate regression splines
- `j` indexes the HLA Class II allotypes `(j = 1, 2, …, J)`
- `I(hla = j)` equals 1 when the observed peptide has HLA allotype `j`, 0 otherwise

There are three levels of random effects:

1. Donor-level effects: `u_donor ∼ Normal(0, σ²_donor)`
2. Donor-protein interaction effects: `u_donor:protein ∼ Normal(0, σ²_donor:protein)`
3. Protein-level effects: `u_protein ∼ Normal(0, σ²_protein)`

Partial pooling via these random effects helps to account for donor heterogeneity and any protein-specific effects.

The variables are:

HLA-II binding scores were estimated using MixMHC2pred and the resulting percent_rank_best scores for each peptide were centred:

- `binding_score_centered`: Centred binding peptide rank score
- `length_diff_from_15`: Difference in peptide length from 15 amino acids
- `hla`: HLA allotype
- `z_sqrt_peptides_donor`: Standardised square root of number of peptides per protein observed per donor
- `z_sqrt_peptides_cohort`: Standardised square root of number peptides peptides per protein observed per cohort
- `donor`: Donor identifier
- `uniprot_id`: UNIPROT protein identifier

Prior Distributions:

- `β`₀ `∼ Normal(-2, 1)` Intercept
- `β`₁ `∼ Normal(-0.25, 0.5)` length_diff_from_15
- `β₁ⱼ ∼ Normal(0, 0.5)` length_diff_from_15:hla interactions
- `β`₂ `∼ Normal(0.5, 0.7)` z_sqrt_peptides_donor
- `β`₃ `∼ Normal(-0.5, 0.7)` z_sqrt_peptides_cohort
- `sⱼ(binding_score_centered) ∼ Normal(0, 1)` Smooth spline prior (sds)
- `σ_donor ∼ Exponential(2)` Donor random effect SD
- `σ_donor:uniprot_id ∼ Exponential(2)` Donor-protein random effect SD
- `σ_uniprot_id ∼ Exponential(2)` Protein random effect SD

## Supporting information

Supplementary information for Immunopeptidomics-guided identification of functional neoantigens in non-small cell lung cancer

Supplementary Data S1: NSCLC Patient Information

Supplementary Data S2: NSCLC Protein-Affecting Variants

Supplementary Data S3: NSCLC Missense Variants

Supplementary Data S4: NSCLC HLA Loss of Heterozygosity

Supplementary Data S5: NSCLC Shared Protein Lists

Supplementary Data S6: NSCLC pVACseq Class I Predictions

Supplementary Data S7: NSCLC pVACseq Class II Predictions

Supplementary Data S8: NSCLC pVACseq Peptidome Combined Predictions

Supplementary Data S9: NSCLC Tested Neoantigens

## Author Contributions

Conceptualisation by PS, TE, CO, PJ, BN and AB. Project administration by PS, TE, CO, PJ and KM. Supervision by PS, TE and CO. Methodology, investigation, visualisation and formal analysis performed by BN, AB, KM, OW and EC. AB, BN, PJ, TE and CO wrote the manuscript. All authors reviewed the manuscript.

## Data availability

Whole Exome Sequencing and RNAseq data have been deposited at the European Genome-phenome Archive (EGA) under EGA Study ID: EGAS00001005499

The mass spectrometry proteomics data have been deposited to the ProteomeXchange Consortium via the PRIDE partner repository with the dataset identifier PXD028990 and 10.6019/PXD028990.

Supplementary Data are available on Github at https://github.com/ab604/lung-neoantigen-supplement and Zenodo https://zenodo.org/doi/10.5281/zenodo.12820423

## References

1. Cancer survival in england - office for national statistics.

2. Cancer Survival in England, cancers diagnosed 2016 to 2020, followed up to 2021.

3. Gainor, J. F. et al. EGFR Mutations and ALK Rearrangements Are Associated with Low Response Rates to PD-1 Pathway Blockade in NonSmall Cell Lung Cancer: A Retrospective Analysis. Clinical Cancer Research 22, 4585–4593 (2016).

4. Mazieres, J. et al. Efficacy of immune-checkpoint inhibitors (ICI) in non-small cell lung cancer (NSCLC) patients harboring activating molecular alterations (ImmunoTarget). Journal of Clinical Oncology 36, 9010–9010 (2018).

5. Jamal-Hanjani, M. et al. Tracking the Evolution of Non–Small-Cell Lung Cancer. New England Journal of Medicine 376, 2109–2121 (2017).

6. Neoantigen-directed immune escape in lung cancer evolution | nature.

7. Bassani-Sternberg, M. et al. Direct identification of clinically relevant neoepitopes presented on native human melanoma tissue by mass spectrometry. Nature Communications 7, 13404 (2016).

8. Nicholas, B. et al. Identification of neoantigens in oesophageal adenocarcinoma. Immunology 168, 420–431 (2023).

9. Newey, A. et al. Immunopeptidomics of colorectal cancer organoids reveals a sparse HLA class I neoantigen landscape and no increase in neoantigens with interferon or MEK-inhibitor treatment. Journal for ImmunoTherapy of Cancer 7, 309 (2019).

10. Marcu, A., et al. The HLA Ligand Atlas - A resource of natural HLA ligands presented on benign tissues. bioRxiv 778944 (2020) doi:10.1101/778944.

11. Andreatta, M. & Nielsen, M. Gapped sequence alignment using artificial neural networks: Application to the MHC class i system. Bioinformatics 32, 511–7 (2016).

12. Juncker, A. S. et al. Systematic Characterisation of Cellular Localisation and Expression Profiles of Proteins Containing MHC Ligands. PLoS ONE 4, e7448 (2009).

13. Müller, M., Gfeller, D., Coukos, G. & Bassani-Sternberg, M. ‘Hotspots’ of antigen presentation revealed by human leukocyte antigen ligandomics for neoantigen prioritization. Frontiers in Immunology 8, (2017).

14. Gfeller, D., Liu, Y. & Racle, J. Contemplating immunopeptidomes to better predict them. Seminars in Immunology 66, 101708 (2023).

15. Wells, D. K. et al. Key Parameters of Tumor Epitope Immunogenicity Revealed Through a Consortium Approach Improve Neoantigen Prediction. Cell 183, 818–834.e13 (2020).

16. Buckley, P. R. et al. Evaluating performance of existing computational models in predicting CD8+ t cell pathogenic epitopes and cancer neoantigens. Briefings in Bioinformatics 23, bbac141 (2022).

17. Gartner, J. J. et al. A machine learning model for ranking candidate HLA class I neoantigens based on known neoepitopes from multiple human tumor types. Nature Cancer 2, 563–574 (2021).

18. Müller, M. et al. Machine learning methods and harmonized datasets improve immunogenic neoantigen prediction. Immunity 56, 2650–2663.e6 (2023).

19. Cai, Y. et al. Immunopeptidomics-guided discovery and characterization of neoantigens for personalized cancer immunotherapy. Science Advances 11, (2025).

20. Huber, F. et al. A comprehensive proteogenomic pipeline for neoantigen discovery to advance personalized cancer immunotherapy. Nature Biotechnology 43, 1360–1372 (2024).

21. Hundal, J. et al. pVACtools: A Computational Toolkit to Identify and Visualize Cancer Neoantigens. Cancer Immunology Research 8, 409–420 (2020).

22. Alexandrov, L. B. et al. Signatures of mutational processes in human cancer. Nature 500, 415–421 (2013).

23. Kandoth, C. et al. Mutational landscape and significance across 12 major cancer types. Nature 502, 333–339 (2013).

24. Gori, K. & Baez-Ortega, A. Sigfit: Flexible Bayesian Inference of Mutational Signatures. 372896 https://www.biorxiv.org/content/10.1101/372896v2 (2020) doi:10.1101/372896.

25. Alexandrov, L. B. et al. The repertoire of mutational signatures in human cancer. Nature 578, 94–101 (2020).

26. Tate, J. G. et al. COSMIC: The catalogue of somatic mutations in cancer. Nucleic Acids Research 47, D941–D947 (2019).

27. Racle, J., Jonge, K. de, Baumgaertner, P., Speiser, D. E. & Gfeller, D. Simultaneous enumeration of cancer and immune cell types from bulk tumor gene expression data. eLife 6, e26476 (2017).

28. Nilsson, J. B. et al. Accurate prediction of HLA class II antigen presentation across all loci using tailored data acquisition and refined machine learning. Science Advances 9, (2023).

29. Jamal-Hanjani, M. et al. Tracking the evolution of nonsmall-cell lung cancer. New England Journal of Medicine 376, 2109–2121 (2017).

30. McGranahan, N. et al. Allele-Specific HLA Loss and Immune Escape in Lung Cancer Evolution. Cell 171, 1259–1271.e11 (2017).

31. Gfeller, D. et al. The Length Distribution and Multiple Specificity of Naturally Presented HLA-I Ligands. The Journal of Immunology (2018) doi:10.4049/jimmunol.1800914.

32. Racle, J. et al. Robust prediction of HLA class II epitopes by deep motif deconvolution of immunopeptidomes. Nature Biotechnology 37, 1283–1286 (2019).

33. Hundal, J. et al. pVAC-seq: A genome-guided in silico approach to identifying tumor neoantigens. Genome Medicine 8, 11 (2016).

34. Gfeller, D. et al. The Length Distribution and Multiple Specificity of Naturally Presented HLA-I Ligands. The Journal of Immunology (2018) doi:10.4049/jimmunol.1800914.

35. Uhlén, M. et al. Tissue-based map of the human proteome. Science 347, (2015).

36. Bürkner, P.-C. brms: An *R* Package for Bayesian Multilevel Models Using *Stan*. Journal of Statistical Software 80, (2017).

37. Howarth, M., Williams, A., Tolstrup, A. B. & Elliott, T. Tapasin enhances MHC class I peptide presentation according to peptide half-life. Proceedings of the National Academy of Sciences 101, 11737–11742 (2004).

38. Boyle, L. H. et al. Tapasin-related protein TAPBPR is an additional component of the MHC class I presentation pathway. Proceedings of the National Academy of Sciences 110, 3465–3470 (2013).

39. Müller, M., Gfeller, D., Coukos, G. & Bassani-Sternberg, M. ‘Hotspots’ of antigen presentation revealed by human leukocyte antigen ligandomics for neoantigen prioritization. Frontiers in Immunology 8, (2017).

40. Bassani-Sternberg, M., Pletscher-Frankild, S., Jensen, L. J. & Mann, M. Mass Spectrometry of Human Leukocyte Antigen Class I Peptidomes Reveals Strong Effects of Protein Abundance and Turnover on Antigen Presentation. Molecular & Cellular Proteomics 14, 658–673 (2015).

41. Gfeller, D. et al. The Length Distribution and Multiple Specificity of Naturally Presented HLA-I Ligands. The Journal of Immunology (2018) doi:10.4049/jimmunol.1800914.

42. Rosenberg, S. A. et al. Immunologic and therapeutic evaluation of a synthetic peptide vaccine for the treatment of patients with metastatic melanoma. Nature Medicine 4, 321–327 (1998).

43. Weinschenk, T. et al. Integrated functional genomics approach for the design of patient-individual antitumor Vaccines1. Cancer Research 62, 5818–5827 (2002).

44. Hsiue, E. H.-C. et al. Targeting a neoantigen derived from a common *TP53* mutation. Science 371, (2021).

45. Castle, J. C. et al. Exploiting the mutanome for tumor vaccination. Cancer Research 72, 1081–1091 (2012).

46. Levine, A. J., Jenkins, N. A. & Copeland, N. G. The Roles of Initiating Truncal Mutations in Human Cancers: The Order of Mutations and Tumor Cell Type Matters. Cancer Cell 35, 10–15 (2019).

47. Hollstein, M., Sidransky, D., Vogelstein, B. & Harris, C. C. p53 Mutations in Human Cancers. Science 253, 49–53 (1991).

48. Lin, M. J. et al. Cancer vaccines: the next immunotherapy frontier. Nature Cancer 3, 911–926 (2022).

49. Vadakekolathu, J. et al. Multi-Omic Analysis of Two Common P53 Mutations: Proteins Regulated by Mutated P53 as Potential Targets for Immunotherapy. Cancers 14, 3975 (2022).

50. Pearson, H. et al. MHC class Iassociated peptides derive from selective regions of the human genome. Journal of Clinical Investigation 126, 4690–4701 (2016).

51. Restrepo-Pérez, L., Joo, C. & Dekker, C. Paving the way to single-molecule protein sequencing. Nature Nanotechnology 13, 786–796 (2018).

52. Alfaro, J. A. et al. The emerging landscape of single-molecule protein sequencing technologies. Nature Methods 18, 604–617 (2021).

53. Lucas, F. L. R., Versloot, R. C. A., Yakovlieva, L., Walvoort, M. T. C. & Maglia, G. Protein identification by nanopore peptide profiling. Nature Communications 12, 5795 (2021).

54. Motone, K. & Nivala, J. Not if but when nanopore protein sequencing meets single-cell proteomics. Nature Methods 20, 336–338 (2023).

55. Martin-Baniandres, P. et al. Enzyme-less nanopore detection of post-translational modifications within long polypeptides. Nature Nanotechnology 18, 1335–1340 (2023).

56. Motone, K., et al. Multi-pass, single-molecule nanopore reading of long protein strands with single-amino acid sensitivity. (2023).

57. Laumont, C. M. et al. Noncoding regions are the main source of targetable tumor-specific antigens. Science Translational Medicine 10, (2018).

58. Chong, C. et al. Integrated proteogenomic deep sequencing and analytics accurately identify non-canonical peptides in tumor immunopeptidomes. Nature Communications 11, (2020).

59. Ruiz Cuevas, M. V., et al. Most non-canonical proteins uniquely populate the proteome or immunopeptidome. Cell Reports 34, 108815 (2021).

60. Jeck, W. R. et al. Circular RNAs are abundant, conserved, and associated with ALU repeats. RNA 19, 141–157 (2012).

61. Horokhovskyi, Y. et al. An Automated Workflow to Address Proteome Complexity and the Large Search Space Problem in Proteomics and HLA-I Immunopeptidomics. Molecular & Cellular Proteomics 24, 101039 (2025).

62. Ferreira, H. J. et al. Immunopeptidomics-based identification of naturally presented non-canonical circRNA-derived peptides. Nature Communications 15, (2024).

63. Ouspenskaia, T. et al. Unannotated proteins expand the MHC-I-restricted immunopeptidome in cancer. Nature Biotechnology 40, 209–217 (2021).

64. Ely, Z. A. et al. Pancreatic cancerrestricted cryptic antigens are targets for T cell recognition. Science 388, (2025).

65. Rooney, M. S., Shukla, S. A., Wu, C. J., Getz, G. & Hacohen, N. Molecular and genetic properties of tumors associated with local immune cytolytic activity. Cell 160, 48–61 (2015).

66. Joshi, K. et al. Spatial heterogeneity of the T cell receptor repertoire reflects the mutational landscape in lung cancer. Nature Medicine 25, 1549–1559 (2019).

67. Ghorani, E. et al. The T cell differentiation landscape is shaped by tumour mutations in lung cancer. Nature Cancer 1, 546–561 (2020).

68. Sahin, U. et al. Personalized RNA mutanome vaccines mobilize poly-specific therapeutic immunity against cancer. Nature 547, 222–226 (2017).

69. Hilf, N. et al. Actively personalized vaccination trial for newly diagnosed glioblastoma. Nature 565, 240–245 (2019).

70. Rojas, L. A. et al. Personalized RNA neoantigen vaccines stimulate T cells in pancreatic cancer. Nature 618, 144–150 (2023).

71. Sethna, Z. et al. RNA neoantigen vaccines prime long-lived CD8+ T cells in pancreatic cancer. Nature 639, 1042–1051 (2025).

72. Andreatta, M. & Nielsen, M. Gapped sequence alignment using artificial neural networks: Application to the MHC class i system. Bioinformatics 32, 511–7 (2016).

73. Nielsen, M., Lundegaard, C., Lund, O. & Keşmir, C. The role of the proteasome in generating cytotoxic T-cell epitopes: insights obtained from improved predictions of proteasomal cleavage. Immunogenetics 57, 33–41 (2005).

74. Singh-Jasuja, H., Emmerich, N. P. N. & Rammensee, H.-G. The Tübingen approach: identification, selection, and validation of tumor-associated HLA peptides for cancer therapy. *Cancer Immunology*, Immunotherapy 53, 187–195 (2004).

75. Tokita, S. et al. Identification of immunogenic HLA class I and II neoantigens using surrogate immunopeptidomes. Science Advances 10, (2024).

76. Danecek, P. et al. Twelve years of SAMtools and BCFtools. GigaScience 10, (2021).

77. Picard Toolkit. (Broad Institute, 2019).

78. Genomics in the Cloud [Book].

79. Ross, E. M., Haase, K., Van Loo, P. & Markowetz, F. Allele-specific multi-sample copy number segmentation in ASCAT. Bioinformatics 37, 1909–1911 (2021).

80. Benjamin, D. et al. Calling Somatic SNVs and Indels with Mutect2. bioRxiv 861054 (2019) doi:10.1101/861054.

81. Koboldt, D. C. et al. VarScan 2: Somatic mutation and copy number alteration discovery in cancer by exome sequencing. Genome Research 22, 568–576 (2012).

82. Kim, S. et al. Strelka2: fast and accurate calling of germline and somatic variants. Nature Methods 15, 591–594 (2018).

83. Bonfield, J. K. et al. HTSlib: C library for reading/writing high-throughput sequencing data. GigaScience 10, (2021).

84. McLaren, W. et al. The ensembl variant effect predictor. Genome Biology 17, 122 (2016).

85. Chen, S., Zhou, Y., Chen, Y. & Gu, J. Fastp: An ultra-fast all-in-one FASTQ preprocessor. Bioinformatics 34, i884–i890 (2018).

86. Kim, D., Paggi, J. M., Park, C., Bennett, C. & Salzberg, S. L. Graph-based genome alignment and genotyping with HISAT2 and HISAT-genotype. Nature Biotechnology 37, 907–915 (2019).

87. Kovaka, S. et al. Transcriptome assembly from long-read RNA-seq alignments with StringTie2. Genome Biology 20, 278 (2019).

88. Hundal, J. et al. Accounting for proximal variants improves neoantigen prediction. Nature Genetics 51, 175–179 (2019).

89. R Core Team. R: A Language and Environment for Statistical Computing. (R Foundation for Statistical Computing, Vienna, Austria, 2018).

90. Purcell, A. W., Ramarathinam, S. H. & Ternette, N. Mass spectrometrybased identification of MHC-bound peptides for immunopeptidomics. Nature Protocols 14, 1687–1707 (2019).

91. Bailey, A. et al. Characterization of the class i MHC peptidome resulting from DNCB exposure of HaCaT cells. Toxicological Sciences (2020) doi:10.1093/toxsci/kfaa184.

92. Perez-Riverol, Y. et al. The PRIDE database resources in 2022: a hub for mass spectrometry-based proteomics evidences. Nucleic Acids Research 50, D543–D552 (2021).

93. Zhang, J. et al. PEAKS DB: De novo sequencing assisted database search for sensitive and accurate peptide identification. Molecular & Cellular Proteomics 11, M111010587 (2012).

94. Tran, N. H., Zhang, X., Xin, L., Shan, B. & Li, M. De novo peptide sequencing by deep learning. Proceedings of the National Academy of Sciences of the United States of America (2017) doi:10.1073/pnas.1705691114.

95. Chong, C. et al. High-throughput and Sensitive Immunopeptidomics Platform Reveals Profound Interferonγ-Mediated Remodeling of the Human Leukocyte Antigen (HLA) Ligandome*. Molecular & Cellular Proteomics 17, 533–548 (2018).

96. Wickham, H. et al. Welcome to the tidyverse. Journal of Open Source Software 4, 1686 (2019).

97. Jessen, L. E. PepTools - an r-Package for Making Immunoinformatics Accessible. (2018).

