## Supplementary information for Immunopeptidomics-guided identification of functional neoantigens in non-small cell lung cancer for "Immunopeptidomics-guided identification of functional neoantigens in non-small cell lung cancer"

#### Figure S1: ALYREF protein coverage

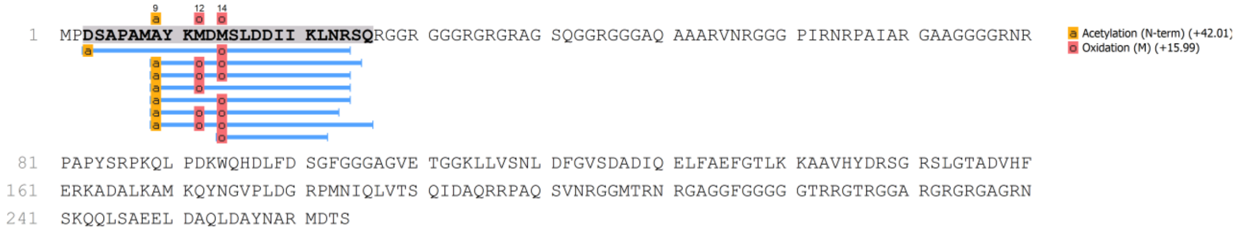


*ALYREF protein coverage table. Nested observed neoantigen peptides are shown mapped in blue*

**Figure S2: MS/MS spectrum of putative neoantigen arising from ALYREF**

A

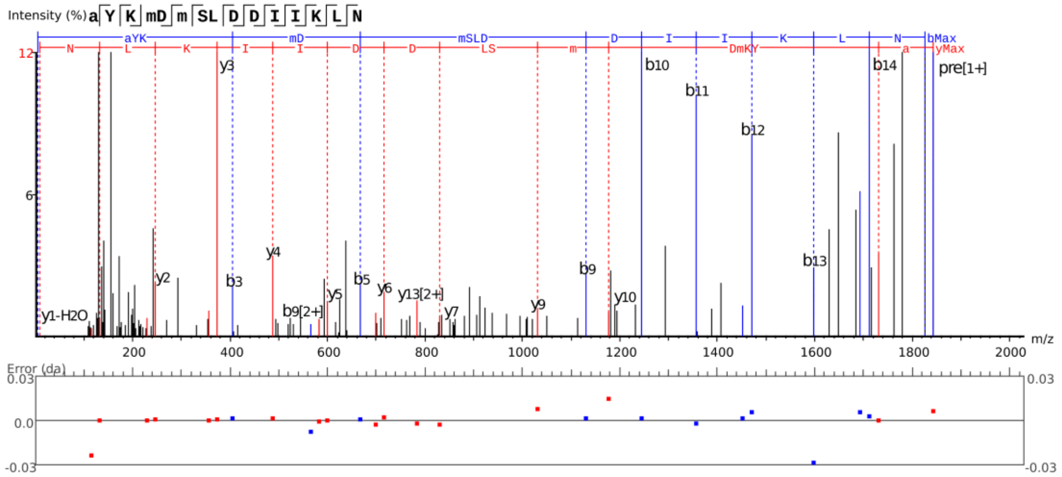


B

| # | b | b-H2O | b-NH3 | b (2+) | Seq | y | y-H2O | y-NH3 | y (2+) | # |
|---|---|---|---|---|---|---|---|---|---|---|
| 1 | 114.1 | 96.0 | 97.0 | 57.5 | A(+42.01) | | | | | 15 |
| 2 | 277.1 | 259.1 | 260.1 | 139.1 | Y | 1730.8 | 1712.8 | 1713.8 | 865.9 | 14 |
| 3 | 405.2 | 387.2 | 388.2 | 203.1 | K | 1567.8 | 1549.8 | 1550.7 | 784.4 | 13 |
| 4 | 552.2 | 534.2 | 535.2 | 276.6 | M(+15.99) | 1439.7 | 1421.7 | 1422.6 | 720.3 | 12 |
| 5 | 667.3 | 649.3 | 650.2 | 334.1 | D | 1292.6 | 1274.6 | 1275.6 | 646.8 | 11 |
| 6 | 814.3 | 796.3 | 797.3 | 407.7 | M(+15.99) | 1177.6 | 1159.6 | 1160.6 | 589.3 | 10 |
| 7 | 901.3 | 883.3 | 884.3 | 451.2 | S | 1030.6 | 1012.6 | 1013.6 | 515.8 | 9 |
| 8 | 1014.4 | 996.4 | 997.4 | 507.7 | L | 943.5 | 925.5 | 926.5 | 472.3 | 8 |
| 9 | 1129.5 | 1111.4 | 1112.4 | 565.2 | D | 830.5 | 812.5 | 813.4 | 415.7 | 7 |
| 10 | 1244.5 | 1226.5 | 1227.5 | 622.7 | D | 715.4 | 697.4 | 698.4 | 358.2 | 6 |
| 11 | 1357.6 | 1339.6 | 1340.5 | 679.3 | I | 600.4 | 582.4 | 583.4 | 300.7 | 5 |
| 12 | 1470.6 | 1452.6 | 1453.6 | 735.8 | I | 487.3 | 469.3 | 470.3 | 244.2 | 4 |
| 13 | 1598.8 | 1580.7 | 1581.7 | 799.9 | K | 374.2 | 356.2 | 357.2 | 187.6 | 3 |
| 14 | 1711.8 | 1693.8 | 1694.8 | 856.4 | L | 246.1 | 228.1 | 229.1 | 123.6 | 2 |
| 15 | | | | | N | 133.1 | 115.1 | 116.0 | 67.0 | 1 |

***(A) MS/MS spectrum of putative neoantigen arising from ALYREF. (B) Ion table.***

**Figure S3: MS/M spectrum of synthetic ALYREF peptide**

A

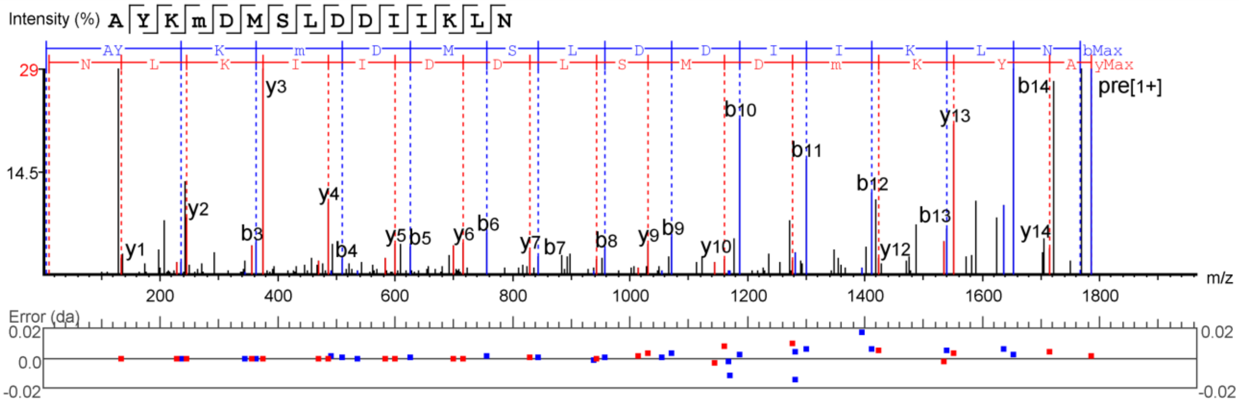


B

| # | b | b-H2O | b-NH3 | b (2+) | Seq | y | y-H2O | y-NH3 | y (2+) | # |
|---|---|---|---|---|---|---|---|---|---|---|
| 1 | 72.0 | 54.0 | 55.0 | 36.5 | A | | | | | 15 |
| 2 | 235.1 | 217.1 | 218.1 | 118.1 | Y | 1714.8 | 1696.8 | 1697.8 | 857.9 | 14 |
| 3 | 363.2 | 345.2 | 346.2 | 182.1 | K | 1551.8 | 1533.8 | 1534.8 | 776.4 | 13 |
| 4 | 510.2 | 492.2 | 493.2 | 255.6 | M(+15.99) | 1423.7 | 1405.7 | 1406.7 | 712.3 | 12 |
| 5 | 625.3 | 607.3 | 608.2 | 313.1 | D | 1276.6 | 1258.6 | 1259.6 | 638.8 | 11 |
| 6 | 756.3 | 738.3 | 739.3 | 378.7 | M | 1161.6 | 1143.6 | 1144.6 | 581.3 | 10 |
| 7 | 843.3 | 825.3 | 826.3 | 422.2 | S | 1030.6 | 1012.6 | 1013.5 | 515.8 | 9 |
| 8 | 956.4 | 938.4 | 939.4 | 478.7 | L | 943.5 | 925.5 | 926.5 | 472.3 | 8 |
| 9 | 1071.4 | 1053.4 | 1054.4 | 536.2 | D | 830.5 | 812.5 | 813.4 | 415.7 | 7 |
| 10 | 1186.5 | 1168.5 | 1169.5 | 593.7 | D | 715.4 | 697.4 | 698.4 | 358.2 | 6 |
| 11 | 1299.6 | 1281.5 | 1282.5 | 650.3 | I | 600.4 | 582.4 | 583.4 | 300.7 | 5 |
| 12 | 1412.6 | 1394.6 | 1395.6 | 706.8 | I | 487.3 | 469.3 | 470.3 | 244.2 | 4 |
| 13 | 1540.7 | 1522.7 | 1523.7 | 770.9 | K | 374.2 | 356.2 | 357.2 | 187.6 | 3 |
| 14 | 1653.8 | 1635.8 | 1636.8 | 827.4 | L | 246.1 | 228.1 | 229.1 | 123.6 | 2 |
| 15 | | | | | N | 133.1 | 115.1 | 116.0 | 67.0 | 1 |

*(A) MS/MS spectrum of synthetic peptide arising from ALYREF. (B) Ion table.*

**Figure S4: LUAD HLA-I model trace plot**

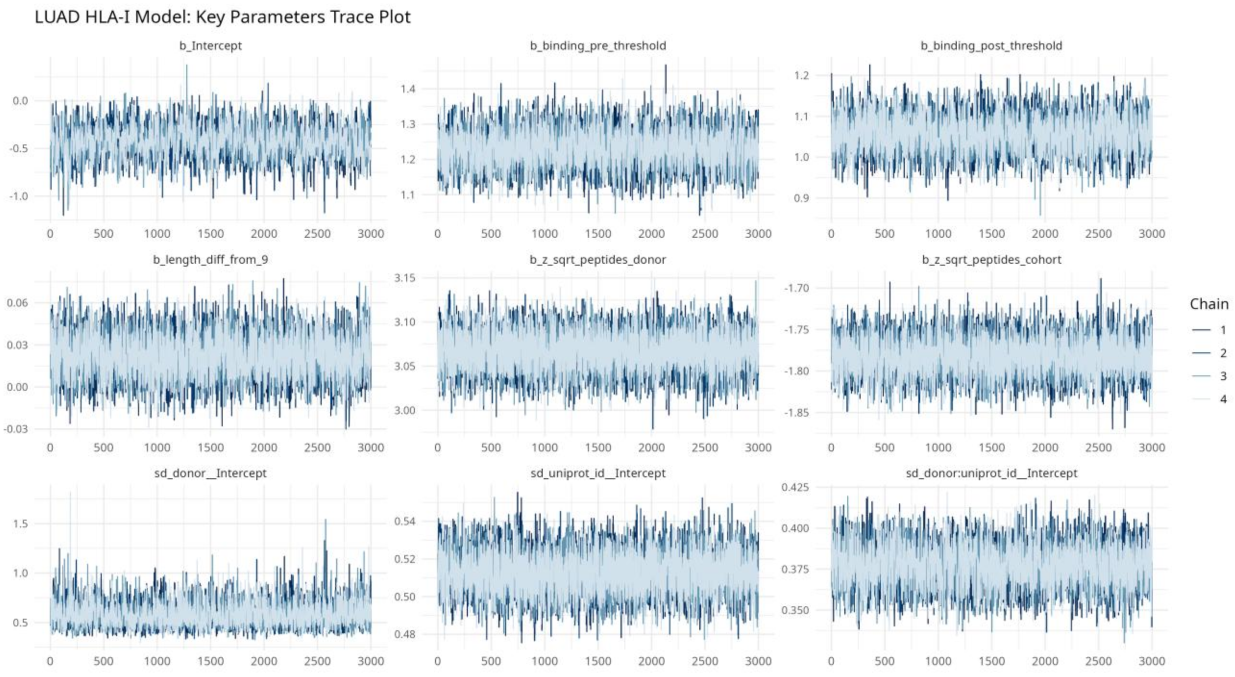


***Trace plot of MCMC draws for key parameters.***

**Figure S5: LUAD HLA-II model trace plot**

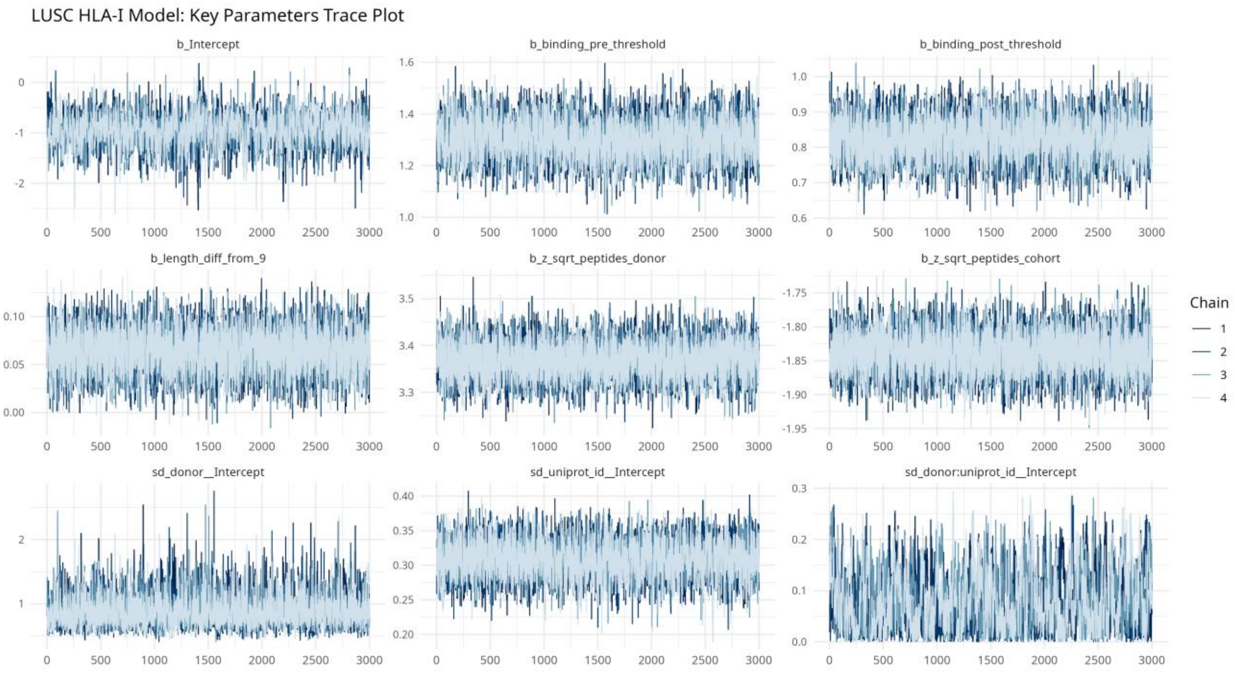


***Trace plot of MCMC draws for key parameters.***

**Figure S6: LUSC HLA-I model trace plot**

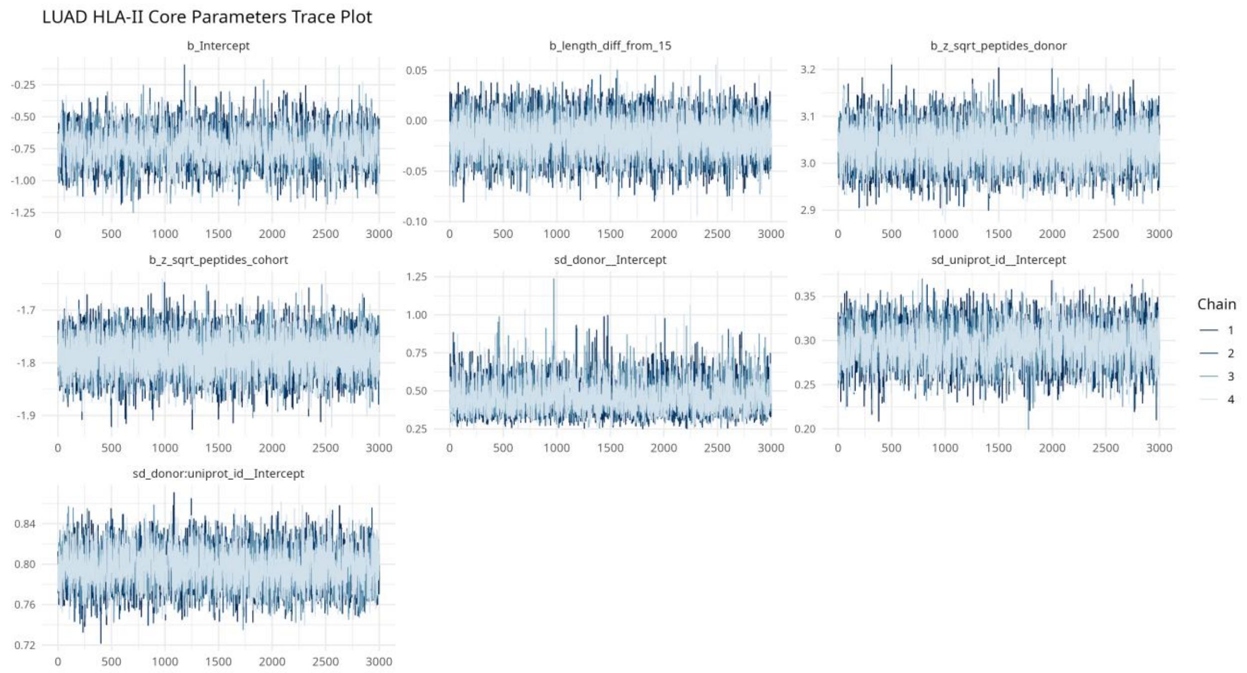


***Trace plot of MCMC draws for key parameters.***

**Figure S7: LUSC HLA-II model trace plot**

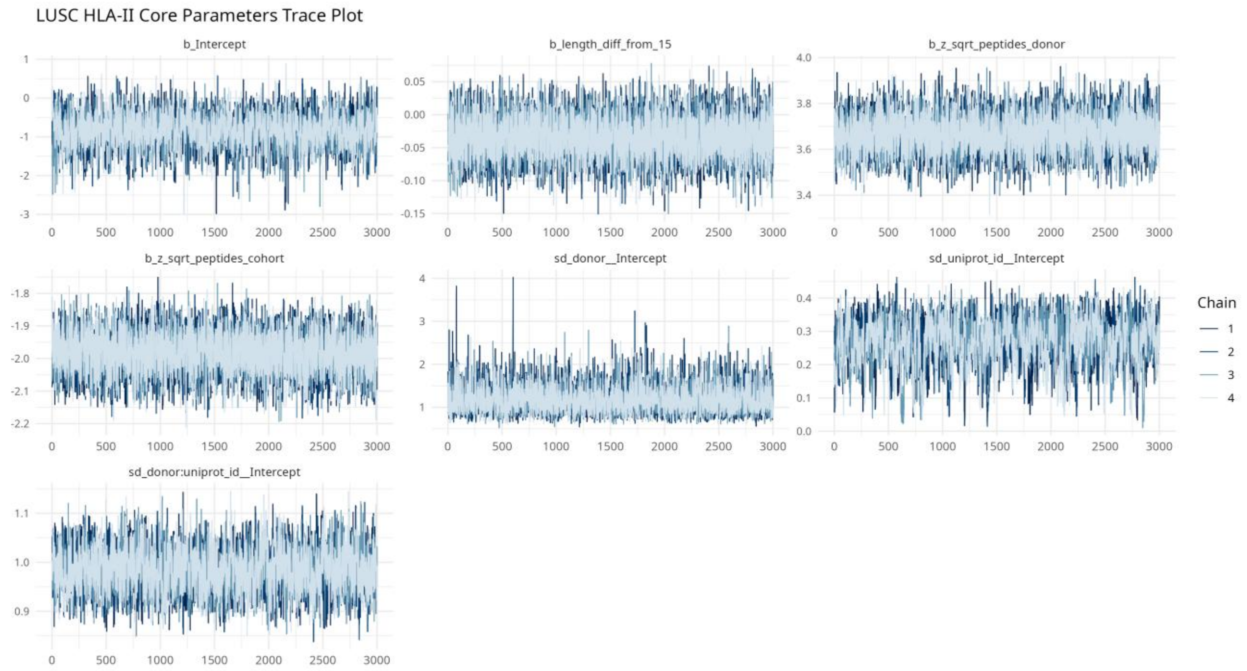


***Trace plot of MCMC draws for key parameters.***

**Figure S8: LUAD HLA-I caterpillar plot**

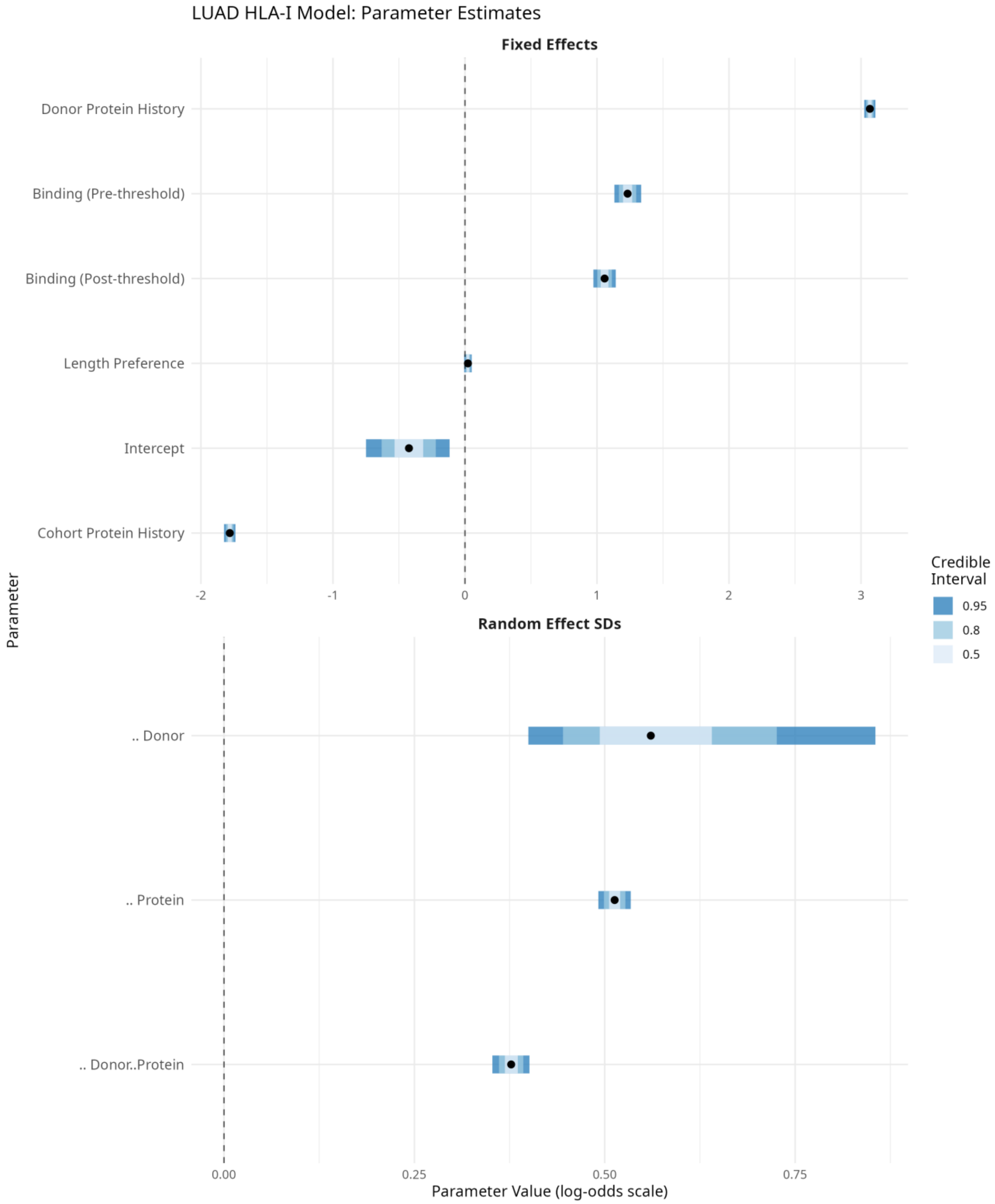


***Caterpillar plot of uncertainty for key model parameters.***

**Figure S9: LUAD HLA-II caterpillar plot**

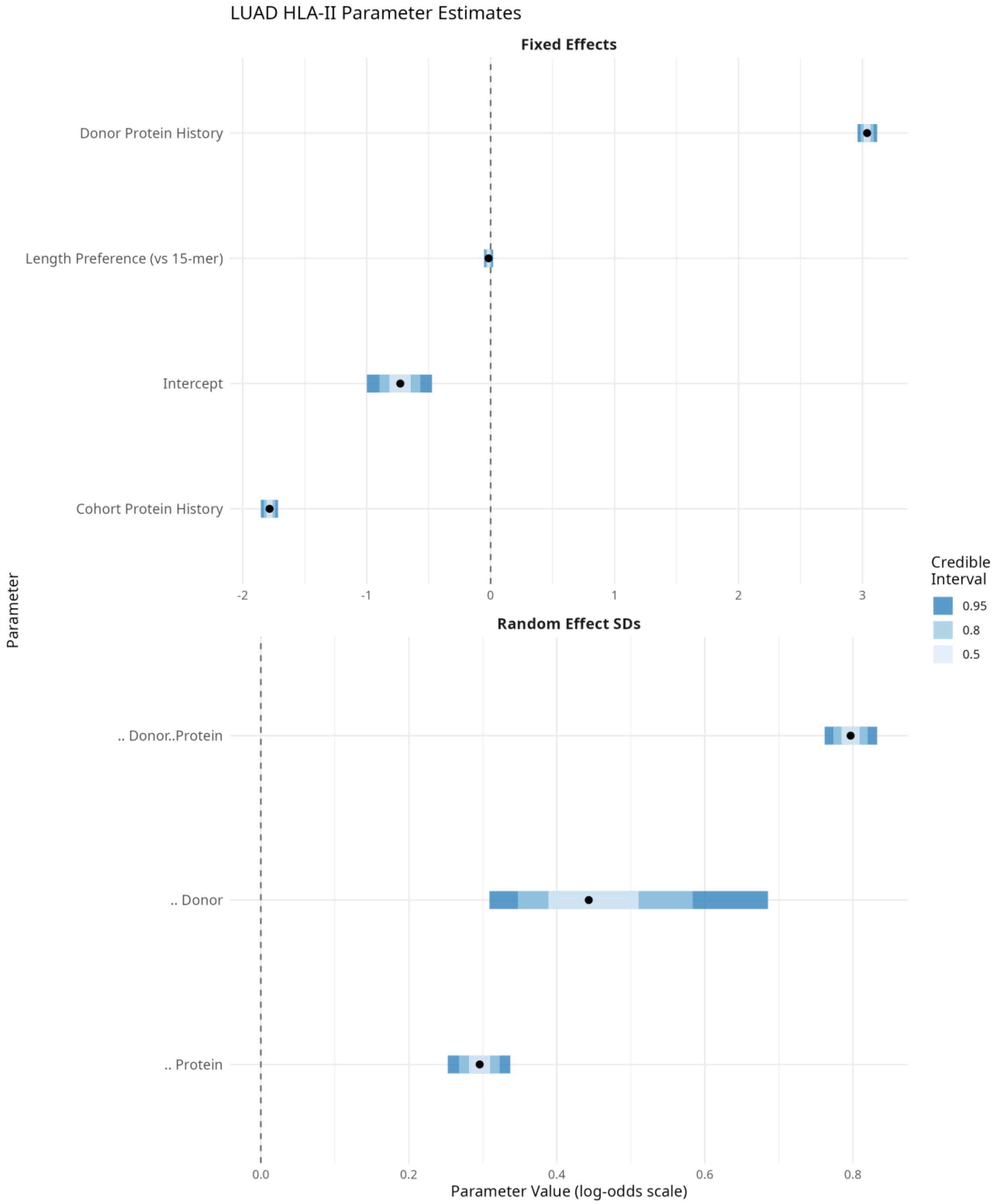


***Caterpillar plot of uncertainty for key model parameters.***

#### Figure S10: LUSC HLA-I caterpillar plot

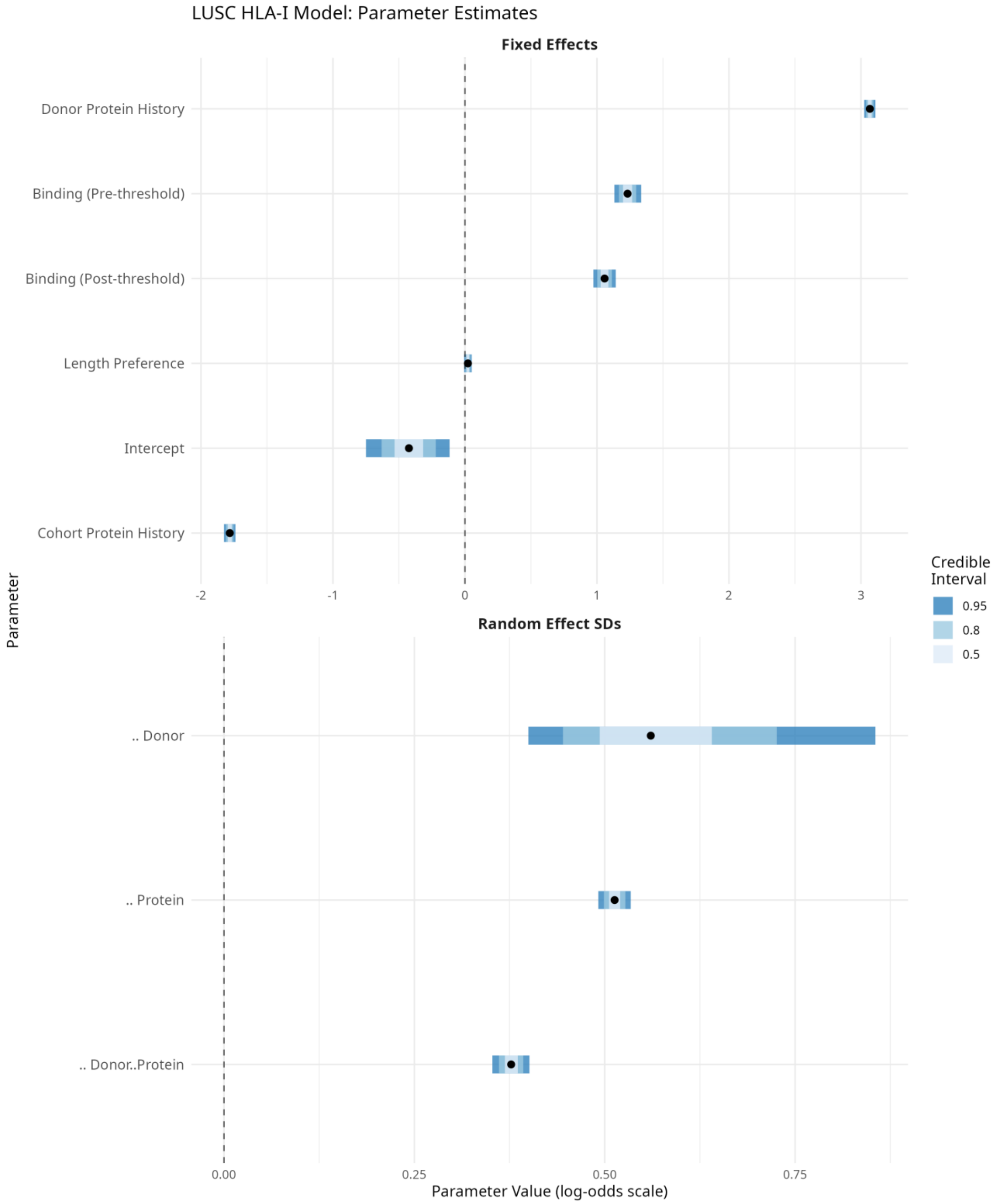


***Caterpillar plot of uncertainty for key model parameters.***

#### Figure S11: LUSC HLA-II caterpillar plot

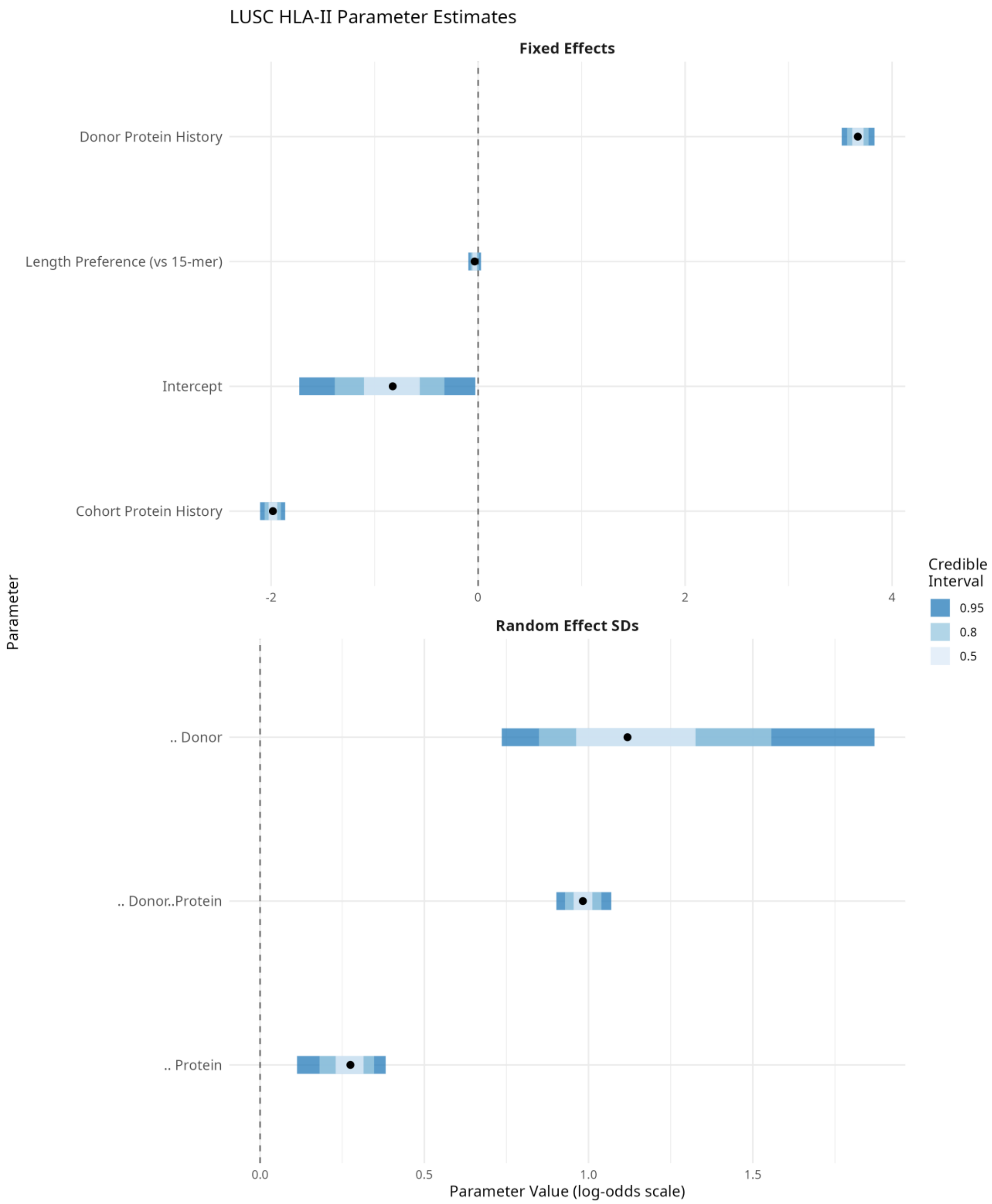


***Caterpillar plot of uncertainty for key model parameters.***

#### Table S1: Bayesian Model Precision Summary

Posterior distributions of key model parameters showing mean, median, standard deviation (sd), 95% credible intervals (ci_lower, ci_upper), probability of direction (pd), proportion of posterior outside ROPE (outside_rope), and signal-to-noise ratio (snr). High SNR values (>10) indicate well-estimated parameters. All parameters show pd = 1, indicating clear directional effects.

***Table S1: Model precision summary***

| model | parameter | mean | median | sd | ci_lower | ci_upper | ci_width | pd | outside_rope | snr |
|---|---|---|---|---|---|---|---|---|---|---|
| LUAD HLA-I | z_sqrt_peptides_donor | 3.07 | 3.07 | 0.02 | 3.02 | 3.11 | 0.08 | 1.00 | 1.00 | 143.05 |
| LUAD HLA-I | z_sqrt_peptides_cohort | -1.78 | -1.78 | 0.02 | -1.83 | -1.74 | 0.09 | 1.00 | 1.00 | 80.87 |
| LUAD HLA-I | binding_pre_threshold | 1.23 | 1.23 | 0.05 | 1.13 | 1.33 | 0.20 | 1.00 | 1.00 | 23.92 |
| LUAD HLA-I | binding_post_threshold | 1.06 | 1.06 | 0.04 | 0.97 | 1.14 | 0.17 | 1.00 | 1.00 | 24.52 |
| LUAD HLA-I | length_diff_from_9 | 0.02 | 0.02 | 0.01 | -0.01 | 0.05 | 0.06 | 0.94 | 0.00 | 1.53 |
| LUSC HLA-I | z_sqrt_peptides_donor | 3.37 | 3.37 | 0.04 | 3.30 | 3.44 | 0.15 | 1.00 | 1.00 | 88.42 |
| LUSC HLA-I | z_sqrt_peptides_cohort | -1.84 | -1.84 | 0.03 | -1.90 | -1.78 | 0.12 | 1.00 | 1.00 | 60.92 |
| LUSC HLA-I | binding_pre_threshold | 1.30 | 1.30 | 0.08 | 1.16 | 1.45 | 0.30 | 1.00 | 1.00 | 17.09 |
| LUSC HLA-I | binding_post_threshold | 0.82 | 0.82 | 0.06 | 0.71 | 0.94 | 0.23 | 1.00 | 1.00 | 14.13 |
| LUSC HLA-I | length_diff_from_9 | 0.06 | 0.06 | 0.02 | 0.02 | 0.11 | 0.09 | 1.00 | 0.00 | 2.91 |
| LUAD HLA-II | z_sqrt_peptides_donor | 3.04 | 3.04 | 0.04 | 2.96 | 3.12 | 0.16 | 1.00 | 1.00 | 74.58 |
| LUAD HLA-II | z_sqrt_peptides_cohort | -1.78 | -1.78 | 0.04 | -1.85 | -1.71 | 0.14 | 1.00 | 1.00 | 49.81 |
| LUSC HLA-II | z_sqrt_peptides_donor | 3.67 | 3.67 | 0.08 | 3.51 | 3.83 | 0.32 | 1.00 | 1.00 | 45.12 |
| LUSC HLA-II | z_sqrt_peptides_cohort | -1.98 | -1.98 | 0.06 | -2.11 | -1.86 | 0.24 | 1.00 | 1.00 | 32.18 |

### Table S2: Bayesian Effective Sample Size Summary

Variance components and effective sample sizes for the hierarchical Bayesian models. Columns: n_don (number of donors), n_prot (number of proteins), n_obs (total observations), avg_don (average observations per donor), avg_prot (average observations per protein), var_don (donor variance component), var_prot (protein variance component), var_don_prot (donor-protein interaction variance), var_res (residual variance), icc_don (intraclass correlation for donors), icc_prot (intraclass correlation for proteins), icc_don_prot (intraclass correlation for donor-protein interaction), de_don (design effect for donors), de_prot (design effect for proteins), n_eff_don (effective sample size for donors), n_eff_prot (effective sample size for proteins).

***Table S2: Effective sample size summary***

| model | n_don | n_prot | n_obs | avg_don | avg_prot | var_don | var_prot | var_don_prot | var_res | icc_don | icc_prot | icc_don_prot | de_don | de_prot | n_eff_don | n_eff_prot |
|---|---|---|---|---|---|---|---|---|---|---|---|---|---|---|---|---|
| LUAD HLA-I | 14 | 9,335 | 164,586 | 11,756 | 18 | 0.33 | 0.26 | 0.14 | 3.29 | 0.08 | 0.07 | 0.04 | 971.74 | 2.09 | 169 | 78,881 |
| LUSC HLA-I | 8 | 7,421 | 50,746 | 6,343 | 7 | 0.80 | 0.10 | 0.01 | 3.29 | 0.19 | 0.02 | 0.00 | 1,206.63 | 1.13 | 42 | 44,798 |
| LUAD HLA-II | 14 | 2,909 | 84,562 | 6,040 | 29 | 0.21 | 0.09 | 0.64 | 3.29 | 0.05 | 0.02 | 0.15 | 299.71 | 1.58 | 282 | 53,508 |
| LUSC HLA-II | 8 | 1,969 | 26,332 | 3,291 | 13 | 1.37 | 0.07 | 0.97 | 3.29 | 0.24 | 0.01 | 0.17 | 791.78 | 1.16 | 33 | 22,767 |

### Supplementary Data S1: NSCLC Patient Information

Supplementary Data S1, a `csv` file containing patient information with 24 rows and 21 column variables. Each row in Supplementary Data S1 represents observations for a single patient. Table 1 provides descriptions of the values contained in each column of Supplementary Data S1.

***Table 1: Patient information Supplementary Data S1 variables***

| Column Variable | Description |
|---|---|
| accel_id | CRUK Accelerator patient identifier |
| target_lung_id | Targeted Lung Health Check patient identifier |
| tissue | NSCLC type: Adenocarcinoma or Squamous cell carcinoma |
| n_somatic_variants | Total number of somatic variants identified by whole exome sequencing |
| mut_burden_per_mb | Mutational burden: mutations per million bases of DNA. Exome target size was 35.7 Mb |
| obs_class_I | Number of observed HLA I peptides by mass spec. immunopeptidomics |
| obs_class_II | Number of observed HLA II peptides by mass spec. immunopeptidomics |
| HLA | Class I and II HLA allotypes identified by genomic sequencing |
| wet_weight | Wet weight of tumour tissue |

| | |
|---|---|
| tumour_purity | Tumour purity as calculated from WES by ASCAT |
| tumour_ploidy | Tumour ploidy as calculated from WES by ASCAT |
| til_status | Tumour infiltrating T-cell status by immunohistochemistry: Low, Moderate, High or NA |
| weeks_post_surgery | Number of weeks since surgery |
| status_as_of_2021_01_19 | Status since 2021-01-19: Alive, Deceased or NA |
| date_of_diagnosis | Date of diagnosis |
| smoking_status | Smoking status |
| notes_2 | Notes about smoking history |

### Supplementary Data S2: NSCLC Protein-Affecting Variants

***Table 2: NSCLC Protein-Affecting Variants Supplementary Data S2 variables***

| Column Variable | Description |
|---|---|
| accel_id | CRUK Accelerator patient identifier |
| vid | Unique variant identifier |
| chrom | Chromosome |
| pos | Genomic coordinate |
| ref | Reference base |
| alt | Variant base |
| type | Variant type: `snv`, `ins`, `del` or `complex`. Single nucleotide variant, insertion, deletion and complex variant respectively |
| gene_name | HGNC gene name |
| ensg | Ensembl gene identifier |
| ensp | Ensembl protein identifier |
| consequence | Variant consequence annotation |
| biotype | Gene biotype filter: protein_coding |
| canonical | Canonical transcript flag |
| vaf | Variant allele frequency |
| depth | Sequencing depth |
| filter | Variant filter status |

| | |
|---|---|
| info | Information field from VCF file |
| format | Format of VCF variable columns |
| sample_1 | Reference sample VCF variable values corresponding with format |
| sample_2 | Tumour sample VCF variable values corresponding with format |
| tissue | Lung tumour tissue type: `Squamous` or `Adenocarcinoma` |

### Supplementary Data S3: NSCLC Missense Variants

***Table 3: NSCLC Missense Variants Supplementary Data S3 variables***

| Column Variable | Description |
|---|---|
| accel_id | CRUK Accelerator patient identifier |
| vid | Unique variant identifier |
| chrom | Chromosome |
| pos | Genomic coordinate |
| ref | Reference base |
| alt | Variant base |
| type | Variant type: `snv`, `ins`, `del` or `complex`. Single nucleotide variant, insertion, deletion and complex variant respectively |
| gene_name | HGNC gene name |
| ensg | Ensembl gene identifier |
| ensp | Ensembl protein identifier |
| vaf | Variant allele frequency |
| depth | Sequencing depth |
| filter | Variant filter status |
| info | Information field from VCF file |
| format | Format of VCF variable columns |
| sample_1 | Reference sample VCF variable values corresponding |

| | |
|---|---|
| | with format |
| sample_2 | Tumour sample VCF variable values corresponding with format |
| tissue | Lung tumour tissue type: `Squamous` or `Adenocarcinoma` |

### Supplementary Data S4: NSCLC HLA Loss of Heterozygosity

***Table 4: NSCLC HLA Loss of Heterozygosity Supplementary Data S4 variables***

| Column Variable | Description |
|---|---|
| accel_id | CRUK Accelerator patient identifier |
| message | HLA LOH detection message |
| HLA_A_type1 | HLA allele type 1: A, B or C |
| HLA_A_type2 | HLA allele type 2: A, B or C |
| LossAllele | HLA allele that was lost |
| KeptAllele | HLA allele that was retained |

### Supplementary Data S5: NSCLC Shared Protein Lists

***Table 5: NSCLC Shared Protein Lists Supplementary Data S5 variables***

| Column Variable | Description |
|---|---|
| hla_class | HLA class (I or II) |
| intersection | Set intersection identifier |
| n_proteins | Number of patients observations of protein in set intersection |
| uniprot_id | UniProt protein identifier |
| gene_name | HGNC gene name |

### Supplementary Data S6: NSCLC pVACseq Class I Predictions

***Table 6: NSCLC pVACseq Class I Predictions Supplementary Data S6 variables***

| Column Variable | Description |
|---|---|
| sample | CRUK Accelerator patient identifier |
| chromosome | The chromosome of this variant |
| start | The start position of this variant in the zero-based, half-open coordinate system |
| stop | The stop position of this variant in the zero-based, half-open coordinate system |
| reference | The reference allele |
| variant | The alt allele |

| | |
|---|---|
| transcript | The Ensembl ID of the affected transcript |
| transcript_support_level | The transcript support level (TSL) of the affected transcript. `NA` if the VCF entry doesn't contain TSL information |
| ensembl_gene_id | The Ensembl ID of the affected gene |
| variant_type | The type of variant. `missense` for missense mutations, `inframe_ins` for inframe insertions, `inframe_del` for inframe deletions, and `FS` for frameshift variants |
| mutation | The amino acid change of this mutation |
| protein_position | The protein position of the mutation |
| gene_name | The Ensembl gene name of the affected gene |
| hgv_sc | The HGVS coding sequence variant name |
| hgv_sp | The HGVS protein sequence variant name |
| hla_allele | The HLA allele for this prediction |
| peptide_length | The peptide length of the epitope |
| sub_peptide_position | The one-based position of the epitope within the protein sequence used to make the prediction |
| mutation_position | The one-based position of the start of the mutation within the epitope sequence. 0 if the start of the mutation is before the epitope |
| mt_epitope_seq | The mutant epitope sequence |
| wt_epitope_seq | The wildtype (reference) epitope sequence at the |

| | |
|---|---|
| | same position in the full protein sequence. NA if there is no wildtype sequence at this position or if more than half of the amino acids of the mutant epitope are mutated |
| best_mt_score_method | Prediction algorithm with the lowest mutant ic50 binding affinity for this epitope |
| best_mt_score | Lowest ic50 binding affinity of all prediction algorithms used |
| corresponding_wt_score | ic50 binding affinity of the wildtype epitope. NA if there is no WT Epitope Seq |
| corresponding_fold_change | Corresponding WT Score / Best MT Score. NA if there is no WT Epitope Seq |
| tumor_dna_depth | Tumor DNA depth at this position. NA if VCF entry does not contain tumor DNA readcount annotation |
| tumor_dna_vaf | Tumor DNA variant allele frequency (VAF) at this position. NA if VCF entry does not contain tumor DNA readcount annotation |
| tumor_rna_depth | Tumor RNA depth at this position. NA if VCF entry does not contain tumor RNA readcount annotation |
| tumor_rna_vaf | Tumor RNA variant allele frequency (VAF) at this position. NA if VCF entry does not contain tumor RNA readcount annotation |
| normal_depth | Normal DNA depth at this position. NA if VCF entry |

| | |
|---|---|
| | does not contain normal DNA readcount annotation |
| normal_vaf | Normal DNA variant allele frequency (VAF) at this position. NA if VCF entry does not contain normal DNA readcount annotation |
| gene_expression | Gene expression value for the annotated gene containing the variant. NA if VCF entry does not contain gene expression annotation |
| transcript_expression | Transcript expression value for the annotated transcript containing the variant. NA if VCF entry does not contain transcript expression annotation |
| median_mt_score | Median ic50 binding affinity of the mutant epitope across all prediction algorithms used |
| median_wt_score | Median ic50 binding affinity of the wildtype epitope across all prediction algorithms used. NA if there is no WT Epitope Seq |
| median_fold_change | Median WT Score / Median MT Score. NA if there is no WT Epitope Seq |
| mh_cflurry_wt_score | MHCflurry ic50 binding affinity for the wildtype epitope |
| mh_cflurry_mt_score | MHCflurry ic50 binding affinity for the mutant epitope |
| mh_cnuggets_i_wt_score | MHCnuggets-I ic50 binding affinity for the wildtype epitope |
| mh_cnuggets_i_mt_score | MHCnuggets-I ic50 binding affinity for the mutant |

| | |
|---|---|
| | epitope |
| net_mhc_wt_score | NetMHC ic50 binding affinity for the wildtype epitope |
| net_mhc_mt_score | NetMHC ic50 binding affinity for the mutant epitope |
| pick_pocket_wt_score | PickPocket ic50 binding affinity for the wildtype epitope |
| pick_pocket_mt_score | PickPocket ic50 binding affinity for the mutant epitope |
| cterm_7mer_gravy_score | Mean hydropathy of last 7 residues on the C-terminus of the peptide |
| max_7mer_gravy_score | Max GRAVY score of any kmer in the amino acid sequence. Used to determine if there are any extremely hydrophobic regions within a longer amino acid sequence |
| difficult_n_terminal_residue | Is N-terminal amino acid a Glutamine, Glutamic acid, or Cysteine? (T/F) |
| c_terminal_cysteine | Is the C-terminal amino acid a Cysteine? (T/F) |
| c_terminal_proline | Is the C-terminal amino acid a Proline? (T/F) |
| cysteine_count | Number of Cysteines in the amino acid sequence. Problematic because they can form disulfide bonds across distant parts of the peptide |
| n_terminal_asparagine | Is the N-terminal amino acid an Asparagine? (T/F) |
| asparagine_proline_bond_count | Number of Asparagine-Proline bonds. Problematic because they can spontaneously cleave the peptide |

| | |
|---|---|
| b_rank | Rank of binding score: 1/median neoantigen binding affinity. Lower is better |
| f_rank | Rank of fold change: the difference in median binding affinity between neoantigen and wildtype peptide (agretopicity). Higher is better |
| m_rank | Ranks of mutant allele expression: the product of gene_expression and tumor_rna_vaf. Higher is better |
| d_rank | Rank of the tumor_dna_vaf. Higher is better |
| score | A score is calculated from the above ranks with the following formula: b_rank + f_rank + (m_rank * 2) + (d_rank/2). Higher is better |
| rank_score | The score converted to a rank, with the best being 1, splitting ties by first. Lower is better |
| rank_percent | The percentage rank score. Lower is better |

### Supplementary Data S7: NSCLC pVACseq Class II Predictions

***Table 7: NSCLC pVACseq Class II Predictions Supplementary Data S7 variables***

| Column Variable | Description |
|---|---|
| sample | CRUK Accelerator patient identifier |
| chromosome | The chromosome of this variant |
| start | The start position of this variant in the zero-based, half-open coordinate system |
| stop | The stop position of this variant in the zero-based, half-open coordinate system |
| reference | The reference allele |
| variant | The alt allele |
| transcript | The Ensembl ID of the affected transcript |
| transcript_support_level | The transcript support level (TSL) of the affected transcript. `NA` if the VCF entry doesn't contain TSL information |
| ensembl_gene_id | The Ensembl ID of the affected gene |
| variant_type | The type of variant. `missense` for missense mutations, `inframe_ins` for inframe insertions, `inframe_del` for inframe deletions, and `FS` for frameshift variants |
| mutation | The amino acid change of this mutation |

| | |
|---|---|
| protein_position | The protein position of the mutation |
| gene_name | The Ensembl gene name of the affected gene |
| hgv_sc | The HGVS coding sequence variant name |
| hgv_sp | The HGVS protein sequence variant name |
| hla_allele | The HLA allele for this prediction |
| peptide_length | The peptide length of the epitope |
| sub_peptide_position | The one-based position of the epitope within the protein sequence used to make the prediction |
| mutation_position | The one-based position of the start of the mutation within the epitope sequence. 0 if the start of the mutation is before the epitope |
| mt_epitope_seq | The mutant epitope sequence |
| wt_epitope_seq | The wildtype (reference) epitope sequence at the same position in the full protein sequence. NA if there is no wildtype sequence at this position or if more than half of the amino acids of the mutant epitope are mutated |
| best_mt_score_method | Prediction algorithm with the lowest mutant ic50 binding affinity for this epitope |
| best_mt_score | Lowest ic50 binding affinity of all prediction algorithms used |
| corresponding_wt_score | ic50 binding affinity of the wildtype epitope. NA if there is no WT Epitope Seq |

| | |
|---|---|
| corresponding_fold_change | Corresponding WT Score / Best MT Score. NA if there is no WT Epitope Seq |
| tumor_dna_depth | Tumor DNA depth at this position. NA if VCF entry does not contain tumor DNA readcount annotation |
| tumor_dna_vaf | Tumor DNA variant allele frequency (VAF) at this position. NA if VCF entry does not contain tumor DNA readcount annotation |
| tumor_rna_depth | Tumor RNA depth at this position. NA if VCF entry does not contain tumor RNA readcount annotation |
| tumor_rna_vaf | Tumor RNA variant allele frequency (VAF) at this position. NA if VCF entry does not contain tumor RNA readcount annotation |
| normal_depth | Normal DNA depth at this position. NA if VCF entry does not contain normal DNA readcount annotation |
| normal_vaf | Normal DNA variant allele frequency (VAF) at this position. NA if VCF entry does not contain normal DNA readcount annotation |
| gene_expression | Gene expression value for the annotated gene containing the variant. NA if VCF entry does not contain gene expression annotation |
| transcript_expression | Transcript expression value for the annotated transcript containing the variant. NA if VCF entry |

| | |
|---|---|
| | does not contain transcript expression annotation |
| median_mt_score | Median ic50 binding affinity of the mutant epitope across all prediction algorithms used |
| median_wt_score | Median ic50 binding affinity of the wildtype epitope across all prediction algorithms used. NA if there is no WT Epitope Seq |
| median_fold_change | Median WT Score / Median MT Score. NA if there is no WT Epitope Seq |
| mh_cnuggets_ii_wt_score | MHCnuggets-II ic50 binding affinity for the wildtype epitope |
| mh_cnuggets_ii_mt_score | MHCnuggets-II ic50 binding affinity for the mutant epitope |
| cterm_7mer_gravy_score | Mean hydropathy of last 7 residues on the C-terminus of the peptide |
| max_7mer_gravy_score | Max GRAVY score of any kmer in the amino acid sequence. Used to determine if there are any extremely hydrophobic regions within a longer amino acid sequence |
| difficult_n_terminal_residue | Is N-terminal amino acid a Glutamine, Glutamic acid, or Cysteine? (T/F) |
| c_terminal_cysteine | Is the C-terminal amino acid a Cysteine? (T/F) |
| c_terminal_proline | Is the C-terminal amino acid a Proline? (T/F) |
| cysteine_count | Number of Cysteines in the amino acid sequence. |

| | |
|---|---|
| | Problematic because they can form disulfide bonds across distant parts of the peptide |
| n_terminal_asparagine | Is the N-terminal amino acid an Asparagine? (T/F) |
| asparagine_proline_bond_count | Number of Asparagine-Proline bonds. Problematic because they can spontaneously cleave the peptide |
| net_mhci_ipan_wt_score | NetMHCIIpan ic50 binding affinity for the wildtype epitope |
| net_mhci_ipan_mt_score | NetMHCIIpan ic50 binding affinity for the mutant epitope |
| n_nalign_wt_score | NNalign ic50 binding affinity for the wildtype epitope |
| n_nalign_mt_score | NNalign ic50 binding affinity for the mutant epitope |
| sm_malign_wt_score | SMMalign ic50 binding affinity for the wildtype epitope |
| sm_malign_mt_score | SMMalign ic50 binding affinity for the mutant epitope |
| b_rank | Rank of binding score: 1/median neoantigen binding affinity. Lower is better |
| f_rank | Rank of fold change: the difference in median binding affinity between neoantigen and wildtype peptide (agretopicity). Higher is better |
| m_rank | Ranks of mutant allele expression: the product of |

| | |
|---|---|
| | gene_expression and tumor_rna_vaf. Higher is better |
| d_rank | Rank of the tumor_dna_vaf. Higher is better |
| score | A score is calculated from the above ranks with the following formula: b_rank + f_rank + (m_rank * 2) + (d_rank/2). Higher is better |
| rank_score | The score converted to a rank, with the best being 1, splitting ties by first. Lower is better |
| rank_percent | The percentage rank score. Lower is better |

### Supplementary Data S8: NSCLC pVACseq Peptidome Combined Predictions

***Table 8: NSCLC pVACseq Peptidome Combined Predictions Supplementary Data S8 variables***

| Column Variable | Description |
| --- | --- |
| sample_id | CRUK Accelerator patient identifier |
| hla_class | HLA class (I or II) |
| table_name | Identifier in the form `accel_id/predicted_hla_allotype/peptide_length` |
| length | Peptide length |
| hla_length_pref1 | HLA allotype preferred peptide length 1 |
| hla_length_pref2 | HLA allotype preferred peptide length 2 |
| gene_name | HGNC gene name |
| mt_epitope_seq | The mutant epitope sequence |
| median_mt_score | Median ic50 binding affinity of the mutant epitope across all prediction algorithms used |
| corresponding_fold_change | `Corresponding WT Score` / `Best MT Score`. `NA` if there is no `WT Epitope Seq` |
| gene_expression | Gene expression value for the annotated gene containing the variant |
| tumor_rna_vaf | Tumor RNA variant allele frequency (VAF) at this |

| | |
|---|---|
| | position |
| tumor_dna_vaf | Tumor DNA variant allele frequency (VAF) at this position |
| rank_percent | The percentage rank score. Lower is better |
| rank_score | The `score` converted to a rank, with the best being 1, splitting ties by first. Lower is better |
| Obs_I | The number of peptides from the source protein observed by mass spectrometry in HLA-I immunopeptidome |
| Obs_II | The number of peptides from the source protein observed by mass spectrometry in HLA-II immunopeptidome |

### Supplementary Data S9: NSCLC Tested Neoantigens

***Table 9: NSCLC Tested Neoantigens Supplementary Data S9 variables***

| Column Variable | Description |
|---|---|
| accel_id | CRUK Accelerator patient identifier |
| gene_name | Gene |
| mt_epitope_seq | Mutated (neoantigen) peptide sequence |
| wt_epitope_seq | Wildtype peptide sequence |
| peptide_length | Peptide length |
| table_name | Identifier in the form `accel_id/predicted_hla_allotype/peptide_length` e.g. `A119/DRB1*04:04/15` |
| mutation | The mutation `From/To` |
| protein_position | Location of the mutation in the source protein, UNIPROT sequence number |
| Obs_I | The number of peptides from the source protein observed by mass spectrometry in HLA-I immunopeptidome |
| Obs_II | The number of peptides from the source protein observed by mass spectrometry in HLA-II immunopeptidome |
| median_mt_score | The median pVACseq predicted binding affinity of the neoantigen peptide |
| median_wt_score | The median pVACseq predicted binding affinity of the wildtype peptide |
| median_fold_change | The ratio between the median neoantigen affinity and wildtype peptide |

| | |
|---|---|
| | affinity |
| rank_percent | The overall rank percentage for the neoantigen from pVACseq for the peptide of that length and HLA allotype |
| mean_sfc_mt | Mean IFN-γ ELISPOT spot forming cells per million cells for the neoantigen peptide |
| mean_sfc_wt | Mean IFN-γ ELISPOT spot forming cells per million cells for the wildtype peptide |
| elispot_response | ELISPOT response category: Strong, Weak or None |

86
